# Structured connectivity in the cerebellum enables noise-resilient pattern separation

**DOI:** 10.1101/2021.11.29.470455

**Authors:** Tri M. Nguyen, Logan A. Thomas, Jeff L. Rhoades, Ilaria Ricchi, Xintong Cindy Yuan, Arlo Sheridan, David G. C. Hildebrand, Jan Funke, Wade G. Regehr, Wei-Chung Allen Lee

**Affiliations:** Department of Neurobiology, Harvard Medical School, Boston, MA, 02115, USA; Program in Neuroscience, Division of Medical Sciences, Graduate School of Arts and Sciences, Harvard University, Cambridge, MA, 02138, USA; HHMI Janelia Research Campus, Ashburn, VA, 20147, USA; F.M. Kirby Neurobiology Center, Boston Children’s Hospital, Harvard Medical School, Boston, MA, 02115, USA

## Abstract

The cerebellum is thought to detect and correct errors between intended and executed commands^1–3^ and is critical for social behaviors, cognition and emotion^4–6^. Computations for motor control must be performed quickly to correct errors in real time and should be sensitive to small differences between patterns for fine error correction while being resilient to noise^7^. Influential theories of cerebellar information processing have largely assumed random network connectivity, which increases the encoding capacity of the network’s first layer^8–13^. However, maximizing encoding capacity reduces resiliency to noise^7^. To understand how neuronal circuits address this fundamental tradeoff, we mapped the feedforward connectivity in the mouse cerebellar cortex using automated large-scale transmission electron microscopy (EM) and convolutional neural network-based image segmentation. We found that both the input and output layers of the circuit exhibit redundant and selective connectivity motifs, which contrast with prevailing models. Numerical simulations suggest these redundant, non-random connectivity motifs increase discriminability of similar input patterns at a minimal cost to the network’s overall encoding capacity. This work reveals how neuronal network structure can balance encoding capacity and redundancy, unveiling principles of biological network architecture with implications for artificial neural network design.

---

Seminal work in the 1960s and 70s^14,15^ provided the basis for influential theories of cerebellar information processing^8,9^. At the input layer, massive numbers of granule cells (GrCs) are hypothesized to randomly sample a smaller number of mossy fiber (MF) inputs, resulting in expansion recoding (**Fig. 1a,b**). Through non-linear activation of the GrCs, expansion recoding projects MF sensory and motor information into a high dimensional representation facilitating pattern separation^10–13,16^ (**Fig. 1c**). At the output layer, each Purkinje cell (PC) integrates tens of thousands of GrC inputs to form appropriate associations^8,9,17^ and are the sole output of the cerebellar cortex (**Fig. 1a,b**). Association learning is hypothesized to occur through linear decoding^17,18^ and reinforcement learning rules^6,19,20^ and may help generate subnetworks of PCs suitable for ensemble learning^21,22^. This network architecture of expansion and re-convergence is also seen in the mushroom body of fly and the electrosensory lobe of electric fish^13,23–27^, suggesting that cerebellar architecture exemplifies a general principle of neuronal circuit organization critical for separating neuronal activity patterns before associative learning. Indeed, these input and output layers are similar to convolutional and linear output layers in artificial deep neural network architectures^28^ (**Extended Data Fig. 1**).

**Figure 1.**
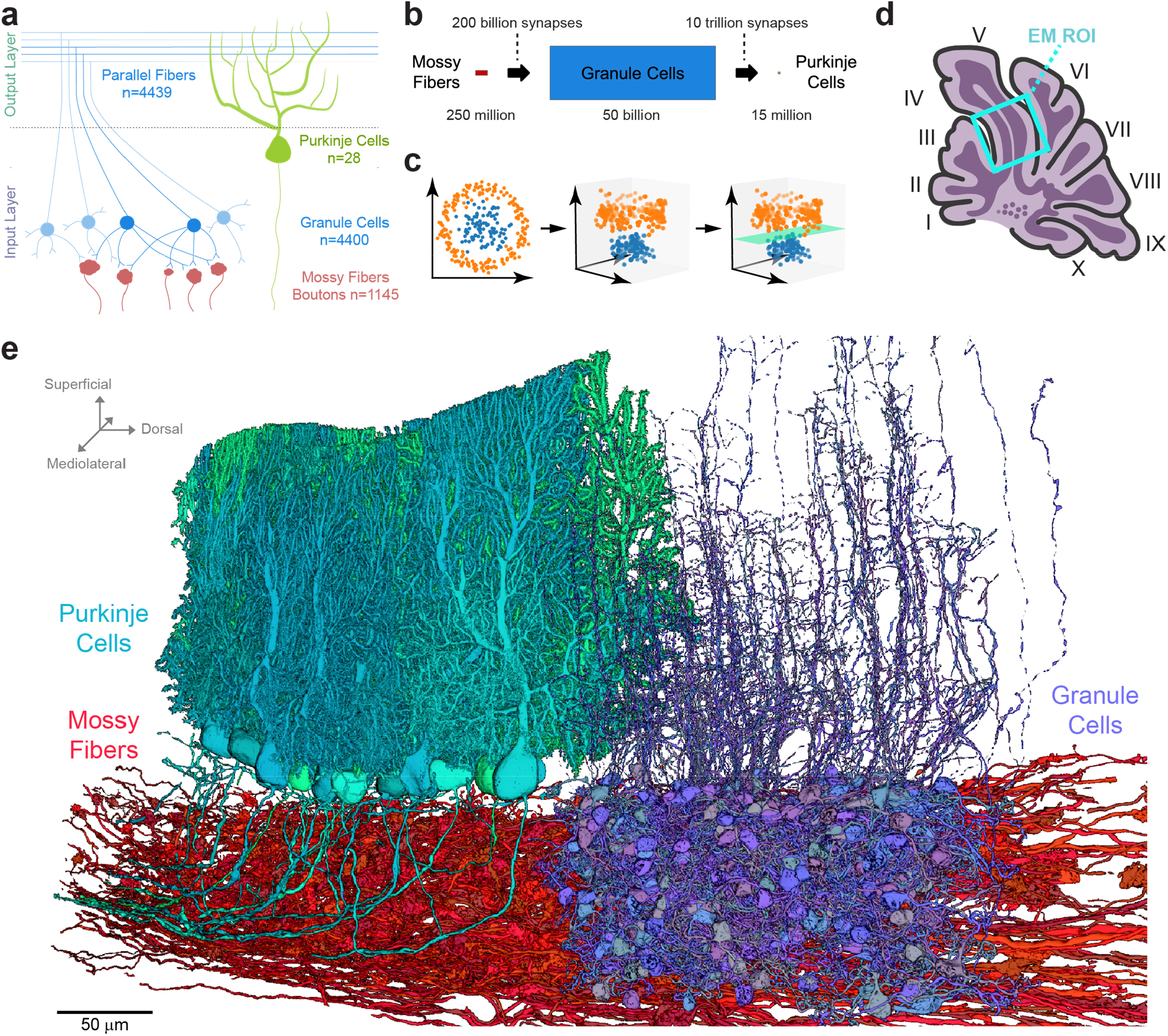
Reconstruction of feedforward circuitry in the cerebellar cortex using large-scale electron microscopy. **a**, Schematic depicting wiring of feedforward neurons in the cerebellar cortex. Granule cells (GrCs, blue circles) sample mossy fiber (MF) boutons (red) and project their axons into the molecular layer where they bifurcate to form parallel fibers. GrC axons make synaptic contacts onto Purkinje cells (PCs, green), which are the sole output of the cerebellar cortex. The number (n) of reconstructed objects (MF boutons and parallel fibers) or cells with cell bodies (GrCs and PCs) in our dataset is shown. **b**, Expansion and convergence of the cerebellar cortex feedforward network. Numbers^46^ denote the global population size (bottom row) and the number of synaptic connections between layers (top row). At the local circuit scale, however, divergence of single MF boutons to GrCs is less (ratio ∼1:3), and convergence of GrCs to PCs is higher (ratio 50,000-200,000:1). **c**, Illustrative data showing how two input representations in 2D (left) once projected into 3D (middle) can be linearly separated (right, green plane). Marr & Albus^8,9^ hypothesize that the MF→GrC dimensionality expansion supports pattern separation and the GrC→PC convergence performs pattern association. **d**, Schematic of a parasagittal section through the vermis of mouse cerebellum with the location of the EM dataset (**Extended Data Fig. 2a**) outlined (cyan box). **e**, 3D rendering of representative EM reconstructions of PCs (green), GrCs (blue) and MFs (red). Non-overlapping GrCs and PCs were rendered for clarity.

Information theoretical models usually assume that MF→GrC connectivity is random in order to maximize the encoding capacity of the input layer of the network^8–11,13^. However, maximizing encoding capacity makes networks less resilient to noise^7^. To understand how the cerebellar cortex balances the tradeoff between encoding capacity and noise resilience, we mapped the precise synaptic connectivity across both input (MF→GrC) and output (GrC→PC) layers of the feedforward circuit in the cerebellar cortex.

We generated a synapse-resolution EM dataset that contains a proximal portion of lobule V from the adult mouse vermis (**Fig. 1d, Extended Data Fig. 2a**). The sample was cut into 1176 serial parasagittal sections, each ∼45 nm thick, and collected onto GridTape^29^, then imaged at a resolution of 4 × 4 nm^2^ per pixel. The dataset was aligned into a 37.2 trillion voxel volume spanning ∼28 million μm^3^ (**Supplementary Video 1**). To facilitate analysis, we densely segmented neurons using artificial neural networks^30^ (**Fig. 1e, Extended Data Fig. 2**). To process a large EM dataset efficiently, we developed a scalable framework to process the dataset in parallel (**Extended Data Fig. 2c**) and a proofreading platform (**Extended Data Fig. 2d**) for targeted neuron reconstruction (**Extended Data Fig. 2e**). We predicted synapse locations and their weights using a separate artificial neural network^31^ trained to identify synapses based on their ultrastructural features (**Extended Data Fig. 2f and 4a,b**) with high accuracy (precision: 95.4%, recall: 92.2%, f-score: 93.8%). In total, we reconstructed >5,500 neurons (**Fig. 1a,e**) and analyzed >150,000 synapses (116,571 MF bouton→GrC and 34,932 GrC→PC) constituting >36,000 unitary connections (13,385 MF bouton→GrC and 23,365 GrC→PC).

We first tested the hypothesis whether connectivity in the input layer (MFs→GrCs) was random. To do so, we densely reconstructed neurons in the GrC layer and identified MFs and GrCs based on morphology^15^. (**Fig. 2a,b, Supplementary Data 1,2, Video 2**). The basic properties of GrCs and MFs were consistent with prior findings^11,32^. GrCs had 4.38 ± 1.00 (mean ± SD) dendrites (**Extended Data Fig. 3a)**, each of which ended in a “claw” that wrapped around and received 10.4 ± 5.6 (mean ± SD) synapses from a single MF bouton (**Extended Data Fig. 4**). GrC dendrites measured 21.7 ± 8.5 μm (mean ± SD) from the cell body center to the center of the presynaptic MF bouton (**Extended Data Fig. 3b**). Individual MF boutons were sampled by 14.8 ± 8.4 (median ± SD) GrCs.

**Figure 2:**
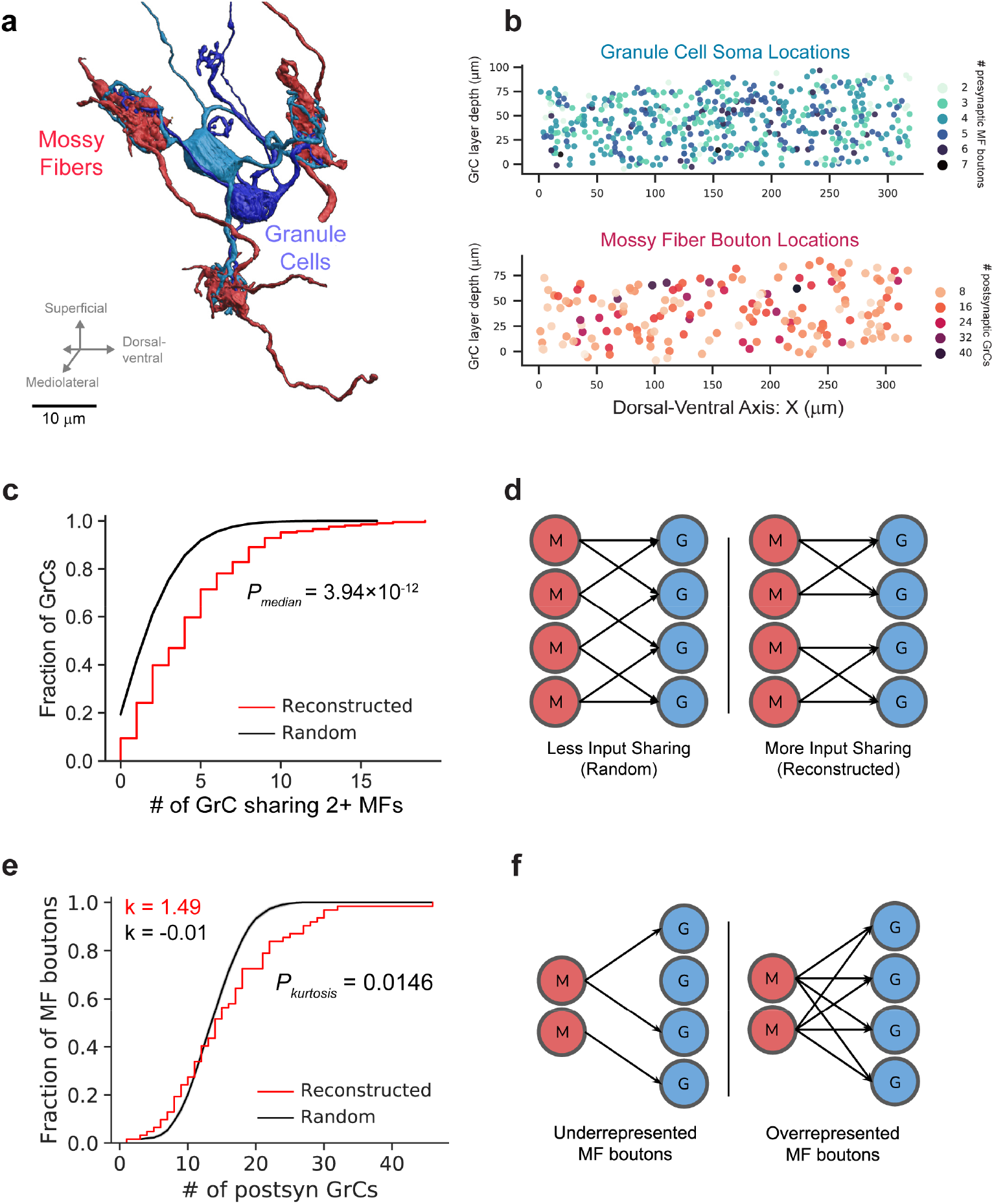
EM reconstructions reveal GrCs redundantly sample MF boutons. **a**, 3D rendering of two GrCs (blue) sharing three common MF bouton inputs. **b**, GrCs (top, n = 4,400) and MF boutons (bottom, n = 1,145) were densely reconstructed. Within a common 320 × 80 × 50 μm subvolume, there are 2,397 GrCs and 784 MF boutons, giving a density of 1,870,000 GrCs and 612,000 MF boutons per mm^3^ and a ratio of 3.06 GrCs per MF bouton. Top: each dot represents a GrC color coded to show the number of dendrites per GrC. Bottom: each dot represents a MF bouton color coded to show the number of postsynaptic GrC targets per bouton. Only neurons in the center 10 μm in Z are plotted for clarity. **c**, Cumulative distribution of MF bouton input redundancy, counting the number of GrC pairs sharing at least 2 MF boutons for each GrC. To minimize edge-effects, only the centermost GrCs are included in this analysis. GrCs in the reconstructed network (red line) share significantly more MF boutons than connectomically-constrained random models (*Radius* model described in **Extended Data Fig. 3d**; also see **Extended Data Fig. 3d-f** for additional random models; black line, shaded regions represent the bootstrapped 95% confidence interval throughout the figures unless otherwise stated; *p* = 3.94 × 10^−12^, n = 211 GrCs, Wilcoxon rank-sum, **Methods**). Note that the reconstructed distribution is plotted as an empirical CDF with discrete numbers of connected neuron pairs, while the random distribution is plotted as CDFs with error shaded across randomly generated networks. **d**, Illustration of redundant sampling in **c** showing pairs of GrCs sharing 2 common MF inputs (right) vs sharing 1 common MF input (left). **e**, Cumulative distribution of postsynaptic GrCs per MF bouton. The reconstructed distribution (red line) is compared with a random model (black line, same as **c**; also see **Extended Data Fig. 3d,l** for other random models). To minimize edge effects, only connections from the centermost MF boutons are counted. Kurtosis (k), a unitless measure of amount of distribution in the tails, is significantly higher in the reconstructed network than the random model suggesting over- and under-sampling of MF boutons by GrCs (*p* = 0.0146, n = 62, Permutation test, **Methods**). **f**, Selective subsampling of MF boutons by the GrCs in **e** creates underrepresented and overrepresented subpopulations.

The Marr-Albus expansion recoding theory predicts random MF→GrC connectivity, which implies minimized sharing of MF bouton inputs between GrCs. In contrast, we found GrC pairs shared 2 or more MF bouton inputs more often than predicted by anatomically-constrained random connectivity^11^ (*p* = 3.9 × 10^−12^, n = 211, Wilcoxon rank-sum test, **Fig. 2c,d and Extended Data Fig. 3d-f**). The distribution of sharing is consistently higher across different numbers of shared inputs (**Extended Data Fig. 3g-i**). Oversharing of inputs among GrCs implies an overconvergence of MF boutons (**Extended Data Fig. 3j)**, and suggests that GrC activity is more correlated than expected, in opposition to the prediction that MF→GrC encoding minimizes population correlations to maximize encoding capacity^7,8,12,13,33^.

Furthermore, we found that MF boutons were not sampled uniformly: the most connected third of MF boutons had 4 times as many postsynaptic GrCs as the least connected third of MF boutons (**Extended Data Fig. 3k**). To rule out the possibility that MF boutons simply had high variability in the number of postsynaptic partners, we quantified the overrepresentation of MF boutons by GrCs with their kurtosis value, a measure of the prevalence of outliers, and found it to be significantly greater than that of random connectivity models (*p*_*kurtosis*_ = 0.0146, n = 62, Permutation test, **Figs. 2e and Extended Data Fig. 3d,l**). By overrepresenting inputs from select MF boutons, these inputs are redundantly encoded by more GrCs, influencing more of the GrC coding space (**Fig. 2f**). Select MF boutons may be more informative or reliable, potentially enhancing noise resiliency of specific inputs. In summary, we found oversharing and overrepresentation to be two redundant wiring motifs that permits expansion coding in the input layer to be more robust to noise and may partially explain low dimensional GrC activity recently reported *in vivo*^34–36^.

In the output layer, given that adjacent PC dendrites share many potential GrC axon inputs (**Fig. 1e and 3a**), we hypothesized that there are redundant and non-random motifs within the GrC→PC connectivity that could contribute to PC synchrony^37–39^. Given the convergence of PCs to downstream DCNs, we propose that structured GrC→PC connectivity can give rise to groups of partially redundant PCs to produce more accurate pattern associations. To test these hypotheses, we densely reconstructed the connectivity between GrC axons and PCs (**Supplementary Data 2-3 and Videos 2-3**). Reconstructed GrC axons were categorized as either “local” or “non-local”. Local GrC axons (n = 541) had cell bodies within the EM volume, while non-local GrC axons (n = 4,439) are parallel fibers without a cell body identified within the dataset (**Fig. 3a**). Dendrites of reconstructed PCs (n = 153 total, n = 28 with somas) ramified across an area of 208 ± 5.2 μm by 158 ± 19.5 μm (mean ± SD) in height and width (**Supplementary Data 3)**, consistent with light microscopy^15^. A common assumption used in computational models is that all GrC-PC pairs that contact each other make synapses^13,17,40,41^. However, we found the connectivity rate was 49 ± 4.4% (mean ± SD) (**Figs. 3a,c and Extended Data Fig. 5a**), with each GrC→PC connection typically comprising one or two synapses (**Extended Data Fig. 4c**), consistent with prior work^42–44^. The lack of all-to-all GrC→PC connectivity raised the question of how selective these unitary connections are and how this selectivity constrains information processing and plasticity mechanisms in the circuit.

**Figure 3:**
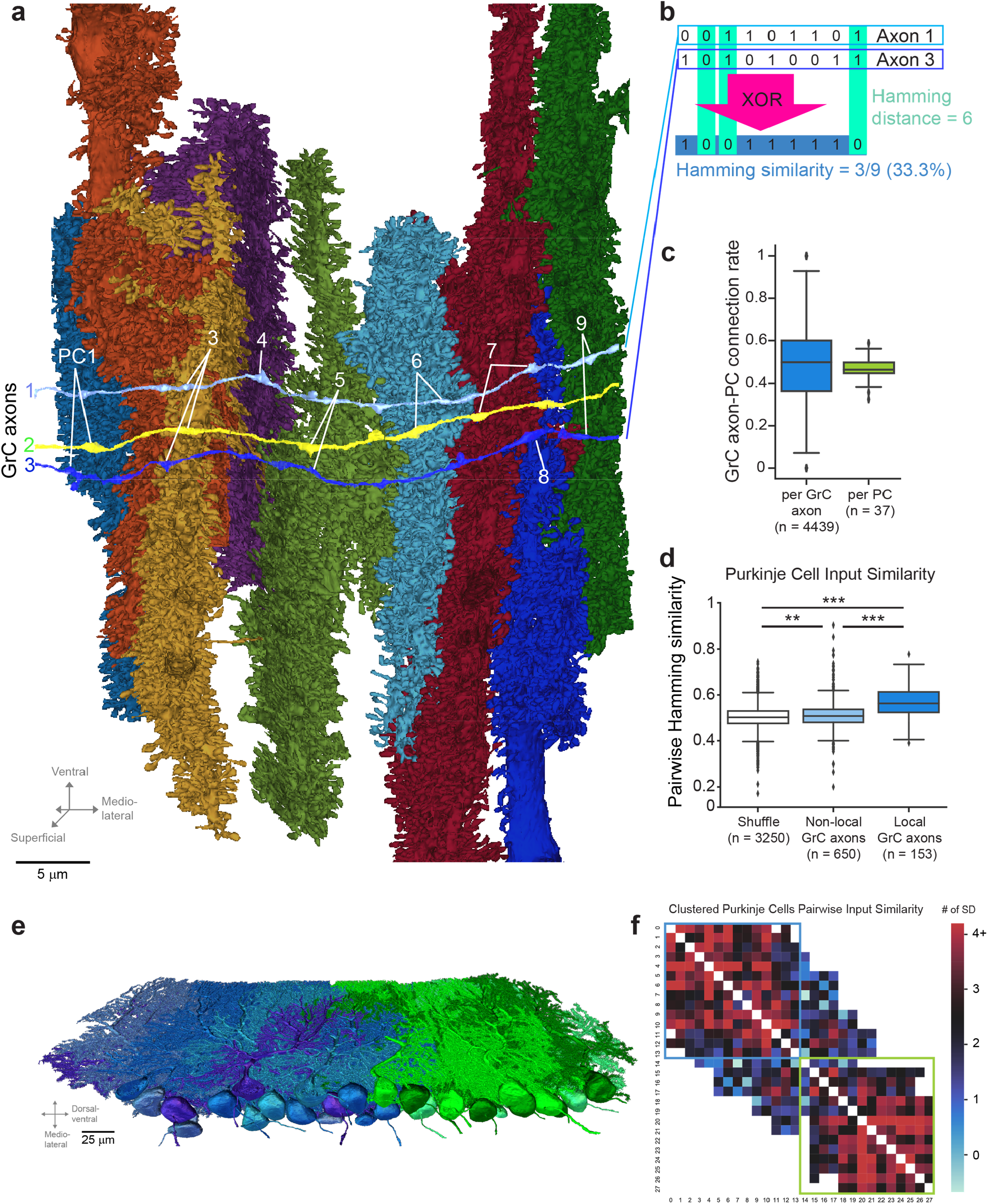
GrC input selectivity predicts PC subnetworks. **a**, 3D EM reconstructions of three GrC axons (colored light blue, yellow, and dark blue), nine PCs, and the locations of synapses (white lines) connecting them. Note that GrC axons have varicosities which are presynaptic to non-PC neurons (e.g., molecular layer interneurons). **b**, Calculation of Hamming similarity as a pairwise metric to compare the similarity of two binary patterns. The example compares the connectivity pattern between the light blue and dark blue GrC axons (1 and 3), with the columns representing different PCs. “1” denotes a connection and “0” denotes the lack of connection. Hamming similarity is similar to cosine or dot-product similarity, but also takes into account the similarity of the non-connections. **c**, Box plot of the ratio of GrC→PC synapses to the total number of times a GrC axon and PC pair contact (touch) one another (**Methods**). Left: synapse ratio per GrC. Right: synapse ratio per PC. **d**, Similarity of GrC inputs between pairs of PCs with at least 30 common GrC axon contacts comparing shuffled input connectivity, non-local GrC axons, and local GrC axons. All three populations are significantly different (*p* = 1.25 × 10^−56^, Kruskal-Wallis test; *p* = 0.00433, shuffle vs. non-local GrC axons; *p* = 9.16 × 10^−32^, non-local GrC axons vs. local GrC axons; *p* = 4.91 × 10^−61^, shuffle vs. non-local GrC axons; Dunn’s post hoc tests). **e**, 3D EM reconstructions of PCs are colored (blue and green) from groupings based on the similarity of their GrC inputs in **d** through spectral clustering which first transformed the input similarity matrix into a low-dimensional embedding before *k*-means clustering (n = 28, *k* = 2, **Methods**). **f**, Input similarity matrix of PCs showing the z-scored similarity of inputs between PCs (color bar, reconstructed input similarity minus random mean divided by the SD). PCs grouped by *k*-means clustering (outlined by blue and green boxes; *k* = 2 as in **e**). Since not all PCs have overlapped inputs with all other PCs, off-diagonal white space denotes uncompared PC pairs.

To examine selectivity in GrC→PC connectivity, we used Hamming similarity as a metric to quantify how similar GrC axon input populations were across PCs, which accounts for both connected and non-connected GrC-PC pairs (**Fig. 3b**). Inputs from local and non-local GrC axons were significantly more similar than shuffled controls, with local GrCs having significantly higher similarity than non-local GrC axons (**Fig. 3d**; local GrC axons vs. shuffle, *p* = 4.9 × 10^−61^; non-local GrC axons vs. shuffle, *p* = 0.0043; non-local GrC axons vs. local GrC axons; *p* = 9.1 × 10^−32^; Kruskal-Wallis and Dunn’s *post hoc* tests), suggesting that PCs exhibit input selectivity. We also detected a small yet significant trend suggesting PCs→GrCs connectivity may be influenced by shared MF bouton inputs (**Extended Data Fig. 5b**, *p* = 1.1 × 10^−4^, Kruskal-Wallis test). The difference between local and non-local inputs may be related to the differences in dimensionality recorded from localized GrC populations^35,36^ versus GrC axons in the molecular layer^45^. We then used spectral clustering to split the PCs into groups based on the similarity of their local GrC inputs (**Fig. 3e,f and Extended Data Fig. 5d**; n = 28, k = 2, **Methods**). Clustering by the similarity of their inputs generated spatially coherent PC groups, each having higher similarity scores within groups than across groups. These groups of PCs may provide a basis for ensemble learning^21,22,46^.

Pattern discrimination and association are most challenging when differences between input patterns are small^7^. To test the effect of the observed connectivity motifs on pattern separation performance, we used dimensionality (**Figs. 1c and 4a**) and signal-to-noise ratio (SNR) (**Fig. 4a**) analyses. Dimensionality measures the number of independent variables embedded in a population’s activity, representing the encoding capacity of the network^12,13^ (**Fig. 1c**). Assuming PCs act as linear decoders^17,18^, SNR measures the ease of discriminating similar patterns. We computed SNR with the signal size defined as the maximum linear distance of GrC representations between two input patterns and noise defined as the variation of the modeled binary activity^12,13^ of the GrC population (**Methods**).

**Figure 4:**
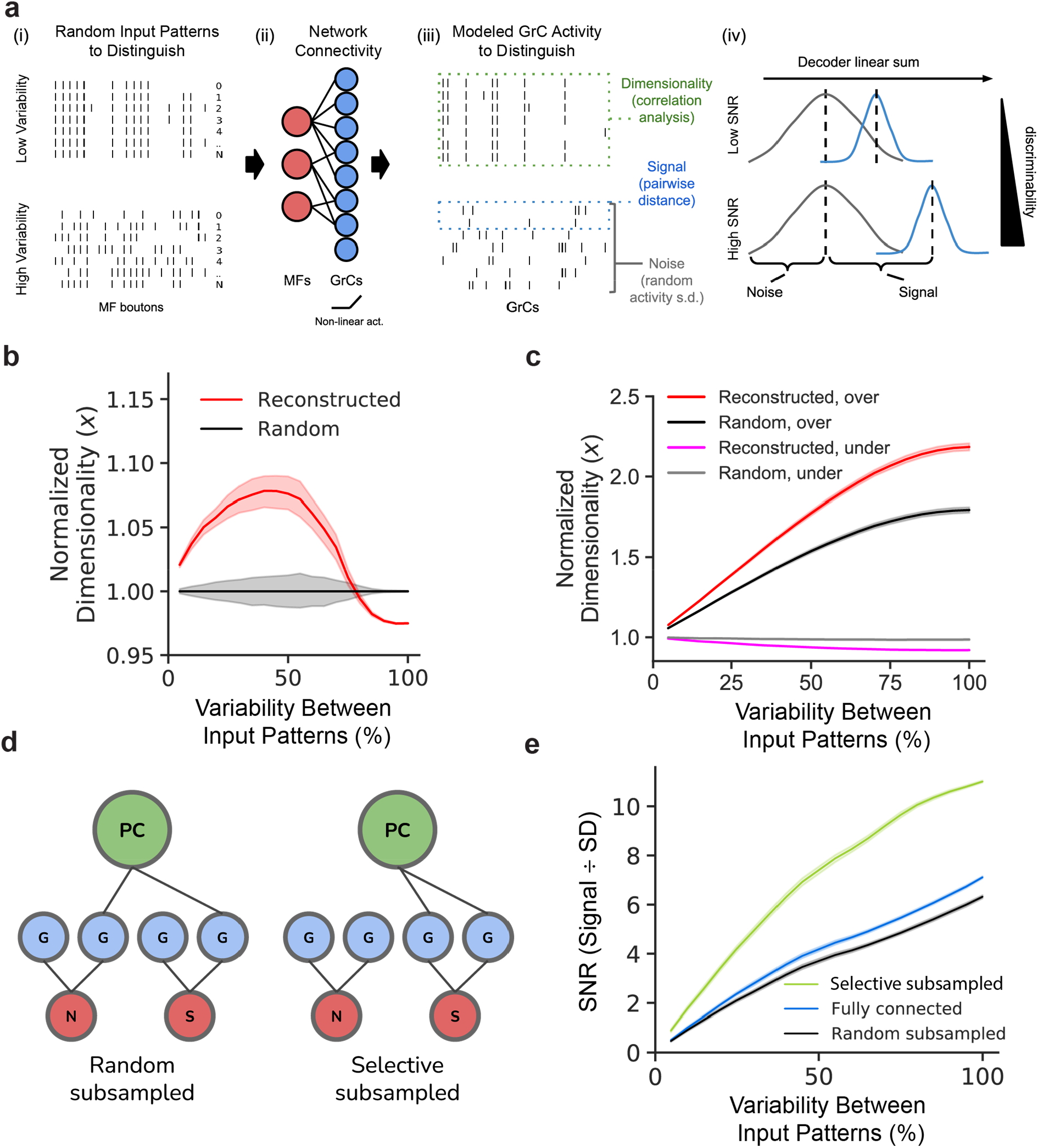
Structured redundancy increases SNR of specific and small input differences. **a**, Dimensionality and signal-to-noise (SNR) analysis. (i) Binary input patterns are modeled with different levels of variability. The objective of the system is to distinguish individual patterns from one another. Note that similar patterns (top, low variability) are harder to distinguish than dissimilar patterns (bottom, high variability). (ii) Input patterns are non-linearly transformed by the MF-GrC network to produce modeled GrC activity^7,13^. (iii) Output activity is analyzed to get dimensionality^13^ (how correlated the activity matrix is), signal (how different/distinguishable each output pattern is from each other), and noise (SD of the linear sum of each pattern). Higher dimensionality and SNR denote better pattern separation performance. (iv) Illustrative histogram of the linear sum of postsynaptic GrC-PC activity. Higher signal relative to noise implies better discriminability. **b**, Relative dimensionality of the GrC population as a function of variability between modeled MF input activity patterns (0% denotes no difference and 100% denotes uncorrelated randomized patterns) comparing the reconstructed (red line) to connectomically-constrained randomly connected models (black line) normalized to the random model. Higher dimensionality indicates higher encoding capacity. The reconstructed network model can encode smaller variability in patterns with greater separability than patterns with larger variations. **c**, Relative dimensionality of the GrC population as a function of variability between MF input patterns, comparing overrepresented (top-third, red and black lines) vs underrepresented (bottom-third, magenta and gray lines) most connected MF boutons in the reconstructed (red and magenta lines) vs random (black and gray lines) connectivity models. Dimensionality is normalized by the underrepresented population in the random connectivity model. **d**, Binary GrC→PC selective subsampling increases SNR. Left: PCs (green) randomly subsample GrCs (blue) regardless of whether they encode a target signal from MFs (red, S) or not (red, N). Right: PCs connect only to GrCs encoding signal-relevant MFs, leading to a higher SNR. **e**, Modeled SNR as a function of variability between input patterns, comparing models with selective (green line), no (blue line), or random (black line) subsampling. Signal size is measured as the maximum difference between postsynaptic linear sums of the modeled input patterns (equivalent to Hamming distance), and noise is defined as the SD of the modeled GrC activity with random MF bouton inputs (**Methods**). Selective sampling enables greater separability by PCs than fully or randomly connected model networks.

At the input layer, we computed the dimensionality of the GrC population in response to different levels of variability in input patterns. Given that random connectivity is thought to maximize dimensionality^7,13^, we predicted that redundant and selective connectivity motifs in the cerebellum would invariably decrease the dimensionality of input patterns. Instead, we found that, when compared to a random connectivity model, the reconstructed network had higher dimensionality for more similar input patterns, but lower dimensionality for largely different input patterns (**Fig. 4b and Extended Data Fig. 6a**). Higher dimensionality for similar input patterns is likely due to an oversharing of MF bouton inputs among GrCs that resulted in input combinations that are more redundant and amplified small input differences, but decreased maximum capacity. Oversharing also increased the convergence rate of MF boutons onto the same sets of GrCs (**Extended Data Fig. 3j)**, increasing the ability to detect coincident changes in those inputs. Selective overrepresentation of inputs (**Fig. 2e,f**) implies that information from specific MF boutons is redundantly encoded by a larger number of GrCs. Overrepresentation of these inputs may help increase SNR (**Fig. 4a**), but should also lower dimensionality due to decreased variation in mixed inputs. Unexpectedly, simulations show that the reconstructed network not only increased SNR (**Extended Data Fig. 6b**), but also expanded dimensionality (**Fig. 4c**), suggesting an increase in capacity to differentiate patterns within the overrepresented MF bouton population (**Fig. 1c**). This may be due to GrCs consistently mixing across over- and under-represented MF bouton populations (**Extended Data Fig. 3k**). However, results also show a substantial decrease to SNR and dimensionality of the underrepresented inputs, suggesting the reconstructed network trade-offs amplifying specific input features over others.

Next, since input subsampling (**Fig. 3c,d**) can result in substantial information loss at the output layer, we sought to understand how selective subsampling can affect decoding from the perspective of PCs. For simplicity, these analyses assumed that GrC inputs were equal in strength. However, we note that some GrC inputs may be more influential than others^47^ (cf.^48–50^) and that >25% of unitary connections have multiple synapses (**Extended Data Fig. 4c**) which would further improve SNR of these inputs. We first measured discriminability by calculating the SNR of the optimal linear distance between patterns and found that while no-subsampling and random-subsampling (**Fig. 4d**) have similar SNRs, selective-subsampling provided a substantial increase in discriminability (**Fig. 4e**), possibly by disconnecting PCs from noisy inputs. Interestingly, while roughly half of potential GrC axon inputs are discarded by subsampling, simulations showed the resulting overall loss in dimensionality was small (∼5% with 50% of GrCs, **Extended Data Fig. 6c**). This suggests a preservation of information across layers by the network, supported by the oversharing and overrepresentation motifs of the input layer.

In summary, at the input layer, we found two redundant connectivity motifs: oversharing of inputs among GrCs and selective overrepresentation of specific MF boutons. Computational analyses suggest that at a small cost to overall capacity, these motifs increase discriminability of similar input patterns, improving robustness of pattern separation despite input variability and synaptic noise^44,51^. These findings run counter to prevailing models of random connectivity where dimensionality is maximized^8–13^ and may partially explain recent findings of lower-than-expected dimensionality of *in vivo* GrC activity^34–36^. Selective overrepresentation of MF boutons, in addition to amplifying signals from more informative inputs to improve robustness to noise, could be a mechanism for differential integration of multimodal MFs^52–54^. Our findings are also compatible with the hypothesis of activity-dependent synaptic plasticity underlying learned representational changes in MF-GrC connectivity^6,55,56^. Future studies incorporating long-range connectivity^53,54,57^ and functional characterization^35,36^ may reveal why some MF boutons are oversampled relative to others.

At the output layer, we found the correlation of GrC inputs among neighboring PCs to be small despite the overlap of their dendritic arbors and correlated spiking patterns^37–39^. Nonetheless, we found clusters of adjacent PCs have higher input similarity, denoting an underlying grouping by function, and suggesting that groups of PCs may be computational units that can be assembled into functional domains and fractured somatotopic maps^58–61^. For individual PCs, this selective connectivity may improve discriminability at the cost of dimensionality. For groups of PCs, decorrelation of anatomical inputs may serve to ensure variability and non-convergent learning, allowing cells in the downstream deep cerebellar nuclei (DCN) to implement ensemble learning, a machine learning approach where many weak and redundant learners (i.e. PCs) are pooled to make a more accurate prediction^21,22,46^.

In conclusion, we examined the feedforward connectivity of the cerebellum at synapse resolution, unveiling the role of non-random redundancy within a circuit that was hypothesized to rely on randomness for maximal performance. Beyond the feedforward network, we expect continued analysis of the dataset will enable the comprehensive examination of cell types making up the cerebellar network^14–16,62–65^. Considering its similarities to modern deep learning architectures (**Extended Data Fig. 1**), our findings of structured connectivity may help to improve artificial intelligence algorithms, build and constrain more complete circuit models, and further our understanding of learning rules^6,19,20^ underlying cerebellar motor control and cognitive function^6,56,66,67^.

## Methods

### Experimental animals

Experimental procedures were approved by the Harvard Medical School Institutional Animal Care and Use Committee and performed in accordance with the Guide for Animal Care and Use of Laboratory Animals and the animal welfare guidelines of the National Institutes of Health. The mouse (*Mus musculus*) used in this study was a ∼P40 male C57BL/6(Jackson Laboratory).

### Sample preparation

The specimen was prepared for EM as described previously^29,68^. We used an enhanced rOTO protocol for heavy metal staining^68,69^ followed by dehydration in a graded ethanol series and embedding in LX112 resin (Ladd Research Industries). The sample was polymerized at 60°C for 2–4 d.

### Sectioning and imaging

The EM dataset of the cerebellum (**Fig. 1d,e and Extended Data Fig. 2a**) was collected and imaged using the GridTape pipeline for automated serial-section TEM^29^. In brief, embedded samples were trimmed (Trim 90, Diatome). Serial ∼45 nm-thick sections were cut using a 35° diamond knife (Diatome) and collected onto LUXFilm-coated GridTape (Luxel Corporation). A total of 1176 sections were cut for a total of ∼49.5 µm total sample thickness. There were 91 single-section losses, 20 instances of adjacent two-section losses, 5 instances of adjacent three-section losses, 2 instances of adjacent four-section losses, and 1 instance of an adjacent five-section loss. Folds, staining artifacts, and cracks occasionally occurred during section processing, but were typically isolated to section edges and therefore not problematic.

Sections were imaged on a JEOL 1200EX transmission electron microscope at 120-kV accelerating potential at 2,500× magnification using a reel-to-reel GridTape stage and a 2 × 2 array of sCMOS cameras (Zyla 4.2, Andor) at 4.3 nm/pixel^29^. Imaging required 521 hours of acquisition on a single tape-based transmission electron microscope with a camera array. The EM images were then stitched together and aligned into a continuous volume using the software AlignTK (https://mmbios.pitt.edu/software#aligntk).

The aligned EM sections were then first imported into CATMAID^70^, then to N5 containers (https://github.com/saalfeldlab/n5) for visualization and segmentation.

### Automated segmentation

We adapted an automated segmentation workflow based on a segmentation pipeline for TEM data^30,71^. In brief, the pipeline consists of three steps: (1) affinity prediction, fragment extraction (over-segmentation), and agglomeration. The affinity prediction uses a 3D U-net convolutional neural network (CNN)^72^ to predict a “connected-ness” probability of each voxel to adjacent voxels. An over-segmentation graph is then produced through a watershed algorithm (2D per-section), and agglomeration is performed iteratively using the “mean affinity” metric^73^ set at 0.5 threshold (segments are only joined when their immediate boundary has affinity greater than 0.5). We modified the original code^71^ (without local shape descriptors) to optimize for runtime performance and to adapt to our supercomputing cluster environment (Harvard Medical School Research Computing). The input to the network is a two-fold downsampled in XY images (effective resolution 8 × 8 × 40 nm); we found the decrease in size and performance to outweigh a minor reduction in accuracy.

#### Affinity CNN training

We trained the CNN using a two-step process: bootstrapped ground truth (GT), and subsequent refinement based on skeleton GT. A set of bootstrapped GT was first created by applying a trained CREMI network (https://cremi.org/) directly on several subvolumes of the dataset, then applying manual corrections using Armitage (Google internal tool). Training on these bootstrapped GT volumes generated an over-segmenting network.

To further improve the accuracy of the network, we scaled up the GT coverage to include 2 regions in the GrC layer, 3 regions in the molecular layer, and 1 region in the Purkinje cell layer – each 6 × 6 × 6 µm (1536 × 1536 × 150 voxels) in size. To accurately and efficiently proofread these volumes, we developed a skeleton-based GT method where we used the connectivity of the manually annotated point-wise skeletons of neurons to correct for errors of the bootstrapped network output. First, human tracers produced dense skeletons of the GT volumes using CATMAID^70^. To correct for split errors, the bootstrapped outputs were agglomerated at very high thresholds (0.7-0.9) to produce an under-segmented graph. Then the GT skeletons are used to correct merge errors, producing an output that can be reviewed and used to improve network training. We iteratively performed the annotation-train-review loop several times per volume as needed (sometimes without further annotation) to converge on a set of high-quality GT volumes.

Using the above GT volumes and supplementary volumes (to be described later), we trained the network using gunpowder (https://github.com/funkey/gunpowder). We used the CREMI network architecture for TEM images^30^. In brief, the U-net had four levels of resolution with downsampling factors in xyz of (3,3,1), (3,3,1), and (3,3,3). The topmost level had 12 feature maps, with the subsequent levels increasing the number of feature maps by 5 fold. Each layer was composed of two convolution passes with kernel sizes of (3,3,3) followed by a rectified linear unit (ReLU) activation function. A final convolution of kernel size (1,1,1) with 3 feature maps and a sigmoid function produced the affinity probability map. The mini batch input size was 268 × 268 × 84 pixels, and the output size was 56 × 56 × 48 pixels. An Adam optimizer was used with learning rate = 0.5 × 10^−4^, beta1 = 0.95, beta2 = 0.999, epsilon = 10^−8^. Gunpowder’s built-in augmentations were used to enable more varied training samples – these include elastic deformation, rotation in the z-axis, artificial slip/shift misalignment, mirroring, transpose, and intensity scale and shift, and missing sections. We also wrote and added duplicated augmentation that randomly selects a section and duplicates it up to 5 times. This is to simulate missing sections where, during deployment, missing sections are replaced with adjacent non-missing data sections. We trained the network to 350,000 iterations.

To evaluate performance of the neural network on unseen data, we further densely traced GT skeletons in another 9 cutouts in various regions of the GrC layer - each spanning 6 × 6 × 6 µm (1536 × 1536 × 150 voxels) in size - and used custom code to evaluate split and merge errors. Compared with other metrics like VOI or RAND index, comparing the number of topological errors is a more direct assessment of the amount of time that proofreaders will take to correct split and merge errors. Plus, producing skeleton GT cutouts is faster, allowing us to make more GT in less time. Evaluating across different artificial neural network parameters, we found that our best segmentation network, at an agglomeration threshold of 0.5, produces an average of 2.33 false merges and 27 false splits per cutout (**Extended Data Fig. 2e)**. Analyzing the results in depth, we found that most false splits belong to small axons at locations with poorly aligned or missing sections.

#### Supplementary ground truth sub-volumes

Beyond the targeted performance evaluation with the nine GT cutouts as described above and before deploying the network across the entire dataset, we had to address problems with segmenting (1) myelin that spans across large regions of the dataset often resulting in merge errors, (2) broken blood vessels that resulted in merge errors with surrounding neurons, (3) darkly stained mitochondria that can make adjacent neuron membranes ambiguous, and (4) GrC axons that are sometimes darkly stained.

To address myelin, we produced 3 additional volumes of regions with dense myelin labeled and trained the network to predict low (i.e., zero) affinity for myelin-containing voxels; the fragment extraction step was then modified to remove fragments that have averaged affinity (average of x, y, z affinity values) less than 0.3. The removal of these fragments reduced myelin-related merge errors.

Blood vessels are relatively large structures in EM that are typically easy to segment. Broken capillaries (during serial sectioning), however, can produce artifacts with misalignments, blending with surrounding neurites and producing merge errors. To address this problem, we used Ilastik^74^ to train a machine learning model (random forest) on 32-fold XY downsampled EM images (effective resolution 128 × 128 × 40 nm) to detect blood vessel voxels and mask them out from the affinity prediction prior to the fragment extraction step.

Mitochondria are often darkly stained in the EM dataset. Because the network is trained to predict high affinities for electron-dense membranous structures including mitochondria, the network can under-predict neuron boundaries when mitochondria are right next to the boundaries creating conditions that can be ambiguous even to human experts. We sparsely annotated an additional GT cutout (3 × 6 × 3 µm in size) with dense mitochondria for training the segmentation network. To our surprise, this worked well without further masking agglomeration or other post processing steps.

Last, there were an infrequent number of GrC axons that were partially darkly stained. These would often be normally stained proximal to the soma but became more darkly stained distally before becoming normally stained again along the ascending portion of the axon. Human annotators can often trace these axons reliably, but they generated errors by the initially trained segmentation network. To mitigate this problem, we made a skeleton GT cutout specifically targeting a region dense with dark GrC axons and added it to the training procedure. Except for regions where multiple darkened axons come together to form a tight bundle of axons, this was effective for segmenting isolated dark axons, speeding up the bulk of the GrC axon reconstruction process.

#### Large-scale parallel processing deployment

To efficiently run the trained CNN and other steps in the automated segmentation pipeline across a large dataset, we developed *daisy* (https://github.com/funkelab/daisy), an open-source work scheduler that can divide the input dataset into spatial chunks and operate on them in parallel (**Extended Data Fig. 2c**). We developed *daisy* with the following design requirements: (1) be scalable and efficient across thousands of workers, (2) be able to tolerate worker and scheduler failures and resumable across reboots, (3) be extensible to non-segmentation tasks, (4) be Python-native for ease of integration with the existing pipelines, and (5) be able to provide on-the-fly dependency calculation between computational stages. At the time of development, no existing tools met such requirements.

#### Targeted neuron reconstruction and proofreading

Although split and merge errors exist in the automated segmentation, such errors are usually easy for humans to recognize in 3D visualizations of reconstructed neurons. Thus, proofreading automated segmentation results is typically faster than manual tracing. That said, merge errors – especially between big neurons and in regions not well-represented by the GT – are endemic for large EM volumes, requiring a low merge-threshold (which creates more split errors), sophisticated un-merge algorithms, and/or exorbitant person-hours to proofread and diagnose where the merge errors happen. These obstacles make using automated large-scale segmentation inaccessible to labs with fewer resources.

To combat the merge error problem, we developed merge-deferred segmentation (*MD-Seg*) proofreading method, a novel workflow that pre-agglomerates fragments only in local blocks (4 × 8 × 8 µm in this work) and defers inter-block merge decisions to proofreaders. In this manner, if there is a merge error, it is limited to the local block and not automatically propagated to hundreds of other blocks which may contain even more merge errors. While deferred, inter-block merge decisions of each neuron are still computed and on-line accessible to proofreaders through a hot-key in the user interface, which is based on Neuroglancer (https://github.com/google/neuroglancer) (**Extended Data Fig. 2d**).

By deferring merge decisions across the dataset, *MD-Seg* allowed us to focus proofreading, targeting regions and cells of interest. Although conventional proofreading can also target specific neurons, because agglomeration is performed globally, a generally high-quality assurance (QA) is needed to be performed for all neurons prior to proofreading to minimize excessive errors among these neurons to the neurons of interest. This pre-proofreading QA burden can be excessive for projects with fewer GT and resources, preventing the start of proofreading. In contrast, in *MD-Seg*, errors between non-targeted neurons do not affect segmentation quality of targeted neurons in most cases. Furthermore, because local blocks are agglomerated separately, it is easier to run segmentation for parts of the dataset, or re-run and update segmentations for specific regions without affecting on-going proofreading progress.

To reconstruct neurons in *MD-Seg*, MFs, GrCs, and PCs were first identified manually based on their stereotypical morphological characteristics^15^ (**Fig. 1f, Supplementary Data 1-3 and Videos 2-3**). Proofreaders typically started by selecting a neuron fragment (contained within a single block), and sequentially ‘grew’ the neuron by adding computed merge decisions of fragments in adjacent blocks through a hot key. During each growth step, the 3D morphology of the neuron was visualized and checked for errors. When merge errors occur, the blocks containing the merge are ‘frozen’ to prevent growth from the merged segment. When a neuron branch stops growing (has no continuations), the proofreader inspects the end of the branch to check for missed continuations (split errors). In this way, both split and merge errors can be corrected.

At any time during growth steps, the proofreader can associate annotations like “uncertain continuations” (potential split errors) as well as periodically save the partial progress of the reconstructed neurons into an on-line database. Upon a saving action, MD-Seg automatically checks if the neuron overlaps with any existing saved neurons, and reports a failure back to the proofreader, enforcing an invariance that any single fragments can only belong to one neuron object. For neurons with higher complexity and overlapping branches (such as a Purkinje cell), to decrease the cognitive load, proofreaders saved the neuron as simpler sub-objects (e.g., purkinje_0.axon, purkinje_0.dendrite_0, …). Once a neuron is marked as completed, a reviewer then loads the neuron to check for possible split or merge errors in a second independent review. Gallery of example reconstructions of MFs, GrCs, and PCs are provided (**Supplementary Data 1-3 and Videos 2-3**)

Using this pipeline, we reconstructed 153 PCs (total; 28 with somas), 732 MF boutons and 2397 GrCs based on morphology. 550 of the 732 MF boutons are from unique MFs, with rare instances of GrCs receiving input from different boutons belonging to the same MF (n = 20). 541 of the 2397 GrCs had axons extending into the molecular layer. We also reconstructed 4439 axons from GrCs whose cell bodies were not present in our volume.

### Automated synapse prediction

We adapted an artificial neural network for synapse prediction on mammalian brain datasets using *synful*^31^ (https://github.com/funkelab/synful). We modified the algorithm to predict synaptic clefts from pointwise ground truth annotations and identified pre- and post-synaptic partners by applying the synapse directionality prediction on the cerebellum dataset. We used webKnossos^75^ to make rough synaptic cleft masks and trained the network to predict these cleft masks. We found that training the network to identify the synaptic clefts produced more generalizable results than either pre- or post-synaptic point predictions and removed ambiguity of one-to-many or many-to-one synapses if only the pre- or post-synaptic partners are predicted. For the network architecture, we used the “small” network size and the cross-entropy loss. We downsampled the GT cutouts by a factor of four in the x- and y-dimension for an effective resolution of 16 × 16 × 40 nm prior to training. Consequently, we reduced the downsampling factors of the U-Net to (2,2,1), (2,2,1), and (2,2,3).

To train and evaluate the synapse prediction network, we annotated nine GT cutouts of the GrC layer (GCL; previously used for evaluating the automated segmentation network, see above), two more of the molecular layer (ML), and one more of the PC layer (PCL) – all of which are 6 × 6 × 6 µm. Five of the GCL cutouts and one of the ML cutouts were used for training – the rest were used for evaluation. Because of the efficiency of pointwise GT generation, we were able to produce a large amount of GT for both training and evaluation. The synapse predictions exhibited excellent accuracy (precision: 95.4%, recall: 92.2%, F-score: 0.938; **Extended Data Fig. 2f**), exceeding that of other TEM datasets^29,31^ (F-score: ∼0.60-0.75) and FIB-SEM datasets^76^ (precision/recall: ∼60-93%). It must be noted, however, that these datasets are of *Drosophila*, which have smaller and more numerous synapses that lead to disagreements even among expert annotators^31,76^. Inspecting the synapse predictions, we found that prediction “errors’’ are typically ambiguous GCL synapses consistent with errors found evaluating synapse predictions elsewhere^76^. In contrast, there were no mispredictions in the ML evaluation dataset where synapses are both larger and clearer in EM (**Extended Data Fig. 4**).

### Quantifying MF→GrC connectivity

We used the reconstructed neurons and the automated synapse predictions to map the connectivity between MF boutons and GrCs. The center of mass of GrCs cell bodies were manually annotated. To accurately get the center of mass of MF boutons, we first collected all MF→GrC synapses (i.e., synapses onto non-GrC neurons were filtered out) associated with a single MF, clustered them into individual boutons through DBSCAN^77^ with parameters eps = 8 µm and min samples = 2, then averaged the synapse locations of each bouton to get the bouton’s center of mass.

#### MF bouton - GrC connectivity proofreading

Beyond optimizing the neural network to minimize false positive (FP) and false negative (FN) rates, we also performed synapse count thresholding and targeted proofreading to more accurately determine binary connectivity between MF boutons and GrCs. Considering that each MF bouton to GrC connection has multiple synapses (**Extended Data Fig. 4c**) and that FPs often would only result in connections with single synapses, simple thresholding would make binary connectivity identification robust even with some FNs. However, we considered two additional factors. First, when MF boutons are at the edge of the volume, only a partial number of synapses are visible, increasing the need for a lower threshold. Second, MF boutons can also seldomly make axon collaterals that make 1 to 2 synapses onto GrC dendrites that are connected to other MF boutons – this is beyond the scope of this paper and so we do not include these collateral synapses in our analyses. Considering these factors, we set the minimum automatic synapse threshold of MF bouton→GrC connections to 3, and then manually validated 2-synapse connections.

#### Counting connectivity spatial distribution

To get the spatial distribution of GrC dendrites in **Extended Data Fig. 3c**, for each GrC, we quantified the spatial displacement (Euclidean distance) of each MF bouton connected to it. To avoid edge effects, we defined margins where we removed from our analysis GrCs within 200 μm of the reconstruction boundaries in the X-axis. We further only counted the GrCs that are either at the top (or bottom) 10 μm of the dataset in Z, removed connections to MFs that are above (or below) the GrC centers of mass in Z, then combined the two distributions. The Y margins do not need to be accounted for since they are natural boundaries. Since the dataset has a Z thickness of 49.5 μm, this method allows unbiased measurement of dendrite distribution of up to 200 μm in X and 39.5 μm in Z. Taken together, we found that GrCs preferentially connect with MF boutons spread in the dorsal-ventral axis rather than in the medial-lateral or the anterior-posterior axis, consistent with previous reports^32^.

#### Counting presynaptic and postsynaptic partners

In **Fig. 2e**, we counted the number of postsynaptic GrCs connected to each MF bouton. To avoid edge effects, we defined margins to not include in the analysis MF boutons within 60 μm and 20 μm to the reconstruction boundaries in X- and Z-axis respectively (Y margins do not need to be accounted for since they are natural boundaries). As shown in **Extended Data Fig. 3c**, these margins capture the extent of the dendritic reach of GrC dendrites. Similarly, for **Extended Data Fig. 3a,b** and **Fig. 2**, we removed analysis GrCs that are within 60 μm and 20 μm of the reconstruction boundaries in X- and Z-axis respectively.

#### MF - GrC random connectivity models

To compare with the observed connectivity in **Fig. 2**, we developed random connectivity models (**Extended Data Fig. 3d**). The “Radius” models are based on random spherical sampling connecting GrCs to MF boutons closest to a single given dendrite length^11–13,32^. “Radius-Average” uses the average dendrite length from our EM reconstructions. The “Radius-Distribution” model is similar, but it uses dendrite lengths drawn from the reconstructed distribution. In the “Vector-Shuffle” model, the dendrite targets of each GrC are drawn (with replacement) from the observed distribution of the spatial displacements between connected GrCs and MF boutons (**Extended Data Fig. 3c**). In all models, the locations of GrCs/MF boutons and the number of dendrites per GrC are maintained to be the same as the reconstructed graph, the margin of searching for MF boutons is set to 10 µm, and there are no double connections between any MF bouton→GrC pair.

### Quantification of GrC→PC connectivity

Similar to MF→GrC connectivity analysis, we used automated segmentation and synapse detection to map GrC→PC connectivity.

#### GrC→PC connectivity proofreading

Since the majority of GrC→PC connections consist of single synapses (**Extended Data Fig. 4c**), it was necessary to proofread GrC→PC synapses to minimize false negative errors. Virtually all false negatives consist of synapses from a GrC axon onto “orphaned” PC spines that are not connected to their PC dendrite. To correct these errors, we found all synapse locations between the reconstructed GrC axons to orphaned segments, and manually proofread them.

#### GrC axon to PC touches

To find locations where GrC axons touch PCs and have the potential, but do not make synapses, we computed and utilized mesh representations of neuron segmentations, where neuron meshes consist of a reduced set of vertices that describe their boundaries. Touches between two neuron meshes were then determined through thresholding the shortest distance between the two sets of vertices; this threshold was set to 10 pixels (at 16 nm mesh resolution) to account for the mesh coordinate approximations. To reduce the number of all-to-all comparisons and improve performance, we simplified vertex locations through downsampling to make rough predictions of distances and prune vertices prior to calculating the exact distances.

#### GrC→PC random connectivity models

To make the “Shuffle” model (i.e., the configuration model) in **Fig. 3d**, for each PC, we shuffled the “connected” status among all connected and touching GrCs preserving the total number of connected GrCs per PC.

#### Computing Hamming similarity

To measure similarity of GrC inputs converging among PCs and outputs of GrCs converging onto PCs, we used Hamming similarity. We used this metric to give connections and the absence of connections equal influence in assessing the similarity of connectivity patterns. Hamming similarity is defined as the inverse of Hamming distance^78^ normalized to the length of the vectors:

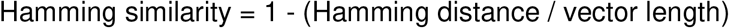

To compare the input convergence of two PCs, for example, that might have different sets of potential presynaptic GrC axons due to differences in the location of their dendrites, we first filtered out all axons that are not either connected to or touching both PCs prior to computing the Hamming similarity score. The same principle was applied to the output convergence of GrC pairs. For **Fig. 3d**, we further filtered out PC pairs that had less than 30 common connected or touching GrC axons to minimize Poisson noise in the analyzed distributions.

#### Purkinje cell clustering

We first calculated a similarity matrix from the PC pairwise Hamming similarity comparing the local GrC input similarity. We then constructed the same matrices for random connectivity GrC→PC models and compared them to the observed matrix to calculate a z-score similarity matrix (**Fig. 3f**). We used spectral clustering^79,80^ to transform the input similarity matrix into a low dimensional embedding prior to performing *k*-means clustering with *k* = 2. For this analysis, we filtered out PC pairs that had less than 10 common contacting GrC axons; the PC pairs that had less than 10 or no common inputs were assigned a z-score of 0. For **Extended Data Fig. 5c**, random clustering was done by taking the computed clusters and randomly shuffling the members (sizes of clusters are preserved). In **Extended Data Fig. 5d**, we performed hierarchical clustering with complete linkage on the same z-score similarity matrix.

### Numerical analyses

We developed a custom framework to generate modeled input and output activity patterns of the MF→GrC layer given the reconstructed (and randomized) MF bouton→GrC connectivity. We assumed the MF input pattern to be binary with a mean activity level of 50%, consistent with prior work^12,13^. Each GrC integrates its inputs based on the reconstructed (or randomized) connectivity graph with equal weights. We assumed a binary output GrC activation function^12,13^, but with an activity level of 30% instead of 10% to be more consistent with recent *in vivo* recordings^35,36^. To maintain a steady-state activity level in a GrC population with a varying number of dendrites per neuron (as observed in **Extended Data Fig. 3a**), we tuned the activation threshold of each GrC by simulating 512 random input patterns and getting the average number of spiking inputs that would produce a 30% activity level for this neuron. Because the MF inputs are binary, however, a single threshold will not produce the precise 30% activity level. As an example, in our configuration, for GrCs with 5 dendrites, an activation threshold of 3 produces a 54% activation rate while a threshold of 4 would produce a 21% activation rate. It is then necessary to further divide the GrC population into an x% population of “low” and a 1-x% population of “high” activation level. In the above example, we solved for *x*% where .21 * *x*% + .54 * (1 - *x*%) = .30, then randomly choose *x*% of GrCs to have an activation threshold of 3 and 1 - *x*% to have 4. Such random assignment of “low” and “high” activation levels allowed us to overcome limitations in previous work^12,13^.

#### Computing dimensionality

To compute the dimensionality of the population activity matrix **x** = (x_0_, x_1_, …) describing activity of the MF or GrC population across trials, we used a previously defined equation^13^:

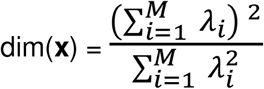

where *λ*_*i*_ are the eigenvalues of the covariance matrix of **x**. Note that this definition is slightly different from the inverse of population correlation^12^, but in general we found the two definitions to give similar results.

For **Fig. 4b,c and Extended Data Fig. 6b**, we constructed the input MF population activity matrix **m** as follows. For each simulation trial, a base vector m_0_ is first randomly generated: each element in m_0_ is drawn independently as a binary number from a Bernoulli distribution with probability 0.5. Derived vectors m_i_ (n = 512 vectors) are then generated from m_0_ with a parameter *x*% (indicating the *x*% difference between input patterns) by randomly rerolling the binary value (not flipped) of *x*% of the vector m_0_. In this manner, vectors created with *x%* = 100% would be entirely uncorrelated to the base input pattern. Each data point (e.g., for each input pattern difference) consists of the average of 100 such simulation trials and the resulting 95% confidence interval.

For **Fig. 4c and Extended Data 6b**, the *x*% parameter (indicating the *x*% difference between input patterns) is restricted and normalized to the 33% MF subpopulation being tested (random, over-, and under-sampled). Within a trial, the variation of MF activity vectors m_i_ is sampled from said fixed MF bouton subpopulations. This is slightly different for **Fig. 4b** where the variation of MF activity vectors m_i_ is sampled across the entire MF population, even if the effective *x*% parameter is the same. For **Extended Data Fig. 6c**, to measure dimensionality of a varying GrC population size with a parameter *y*%, we ran MF**→**GrC simulation as stated, but only performed dimensionality analysis on a random *y*% subset of the GrCs. The subset is re-randomized for each trial.

For **Extended Data Fig. 6a**, we modeled MF inputs as continuous variables representing input spiking frequencies^13^. As for the binary input model, we also performed GrC activation threshold tuning to produce a precise coding level for the GrC population (i.e., 30%), though we did not need “low” and “high” assignments since inputs are continuous. Given a base input vector m_0_, derived input vector m_i_ is derived by adding m_0_ to a random noise vector n_i_ with a noise factor *f*_*noise*_:

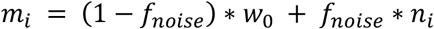

The noise factor *f*_*noise*_ effectively controls the degree of difference between input vectors within the MF activity matrix **m**. At low *f*_*noise*_, **m** is more static, while at high *f*_*noise*_, **m** is composed of more independently drawn random input vectors.

*Computing learned signal size*. It is thought that the GrC population activity is normalized through inhibition from the Golgi cells^8,9^, which is why we tuned the GrC population to have the same mean activity across input MF patterns. Assuming equal weights, the presynaptic sum of GrC activations is expected to be the same across patterns. To differentiate patterns, it is theorized that PCs can manipulate their synaptic weights to affect (increase or decrease) the postsynaptic linear sum of specific GrC activation^7,8,13,17,18,33,40,46^.

For **Fig. 4e**, we defined the learned signal size as the maximum difference between the postsynaptic linear sum of the GrC activations to be distinguished. Given two GrC activation vectors *g*_*j*_ and *g*_*k*_, and a set of weights *w*, the linear sum difference is:

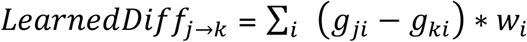

Given the constraints that weights are positive (GrC→PC synapses are excitatory) and that weights range from 0.0 to 1.0, the max learned difference from *g*_*j*_ to *g*_*k*_ would be:

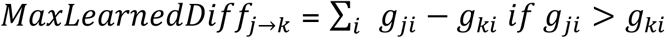

Alternatively, the max learned difference from *g*_*k*_ to *g*_*j*_ is:

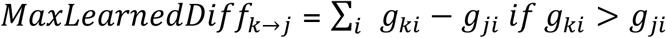

For simplicity and without loss of generality we simply calculated a combined metric, which equates to the Hamming distance between two GrC activation vectors when GrC activation vectors are binary and weights are between 0.0 and 1.0:

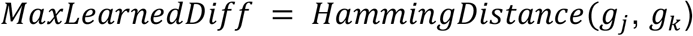

For each simulation trial, we generated a base input MF vector m_0_ and a set of derived vectors *m*_*1*,.., *n*_ (n = 512) as described previously. We then computed the pairwise Hamming distance of the resulting GrC activation vector *g*_*0*_ against *g*_*1, 2*,.., *n*_ to measure the average and variance of responses. Each data point (e.g., for each input pattern difference) consists of 100 such simulation trials and the resulting 95% confidence interval. For the “selective subsampled” model in **Fig. 4e**, we chose the 50% subset of GrCs that would produce the greatest Hamming distance across the derived activation vectors *m*_*1*,.., *n*_ to *m*_*0*_.

#### Computing noise

For **Fig. 4e**, we computed noise as the magnitude of variation of the modeled GrC activity across random background MF input patterns. To calculate the standard deviation of background activity noise, we ran simulations of 512 random MF input patterns and calculated the standard deviation of the sums of GrC outputs.

## Data and Code Availability

Source code for *Daisy* is available at https://github.com/funkelab/daisy. *MD-Seg*’s front-end is available at https://github.com/htem/neuroglancer_pr/tree/segway_pr_v2, and back-end at https://github.com/htem/segway.dahlia. Directions for accessing the acquired EM dataset, reconstructed neuron surface renderings, neuron connectivity graphs, will be available at https://github.com/htem/cb2_project_analysis.

## Acknowledgements

We thank X. Guan, Y. Hu, M. Liu, E. Mayette, M. Narwani, M. Osman, D. Patel, E. Phalen, R. Singh, and K. Yu for neuron reconstruction and manual annotation of ground truth for cell segmentation and synapse predictions; J. Rozowsky for neuron reconstruction, analysis, and the initial finding of redundant MF→GrC connectivity; P. Li and V. Jain for access to and support with Google Armitage/BrainMaps software; Y. Hu and M. Osman for contributing code for segmentation accuracy measurements; M. Narwani, K. Yu and X. Guan for contributing code for the proofreading platform; J. Buhmann and N. Eckstein for discussions and machine learning advice; C. Guo, T. Osorno, S. Rudolph, and L. Witter, for discussion, advice, and help with EM preparations; C. Bolger for EM sample preparation; T. Ayers, R. Smith, and Luxel Corporation for coating GridTape; O. Rhoades for help with illustrations; J. Drugowitsch, A. Handler, C. Ott, R. Wilson, A. Kuan, and members of the Lee lab for comments on the manuscript. This work was supported by the NIH (R21NS085320, RF1MH114047), the Bertarelli Program in Translational Neuroscience and Neuroengineering, Stanley and Theodora Feldberg Fund, and the Edward R. and Anne G. Lefler Center. Portions of this research were conducted on the O2 High Performance Compute Cluster at Harvard Medical School partially provided through NIH NCRR (1S10RR028832-01) and a Foundry Award for the HMS Connectomics Core.

## Author Contributions

T.M.N., L.T., W.R., and W.C.A.L. conceptualized the project and designed experiments. D.G.C.H. sectioned samples. L.T. imaged and aligned the EM dataset. A.S. and J.F. developed the segmentation and synapse prediction networks. T.M.N. and J.F. developed *Daisy*. T.M.N. developed the *MD-Seg* proofreading platform. T.M.N. and L.T. developed and applied the segmentation pipeline. T.M.N., J.L.R., I.R., and X.C.Y. developed and applied the synapse prediction pipeline. T.M.N., L.T., J.L.R., I.R. and X.C.Y. performed reconstructions and analyzed the data. T.M.N., L.T., J.L.R., W.G.R., and W.C.A.L. wrote the paper with input from the other authors.

## Extended Data for

**Extended Data Figure 1.**
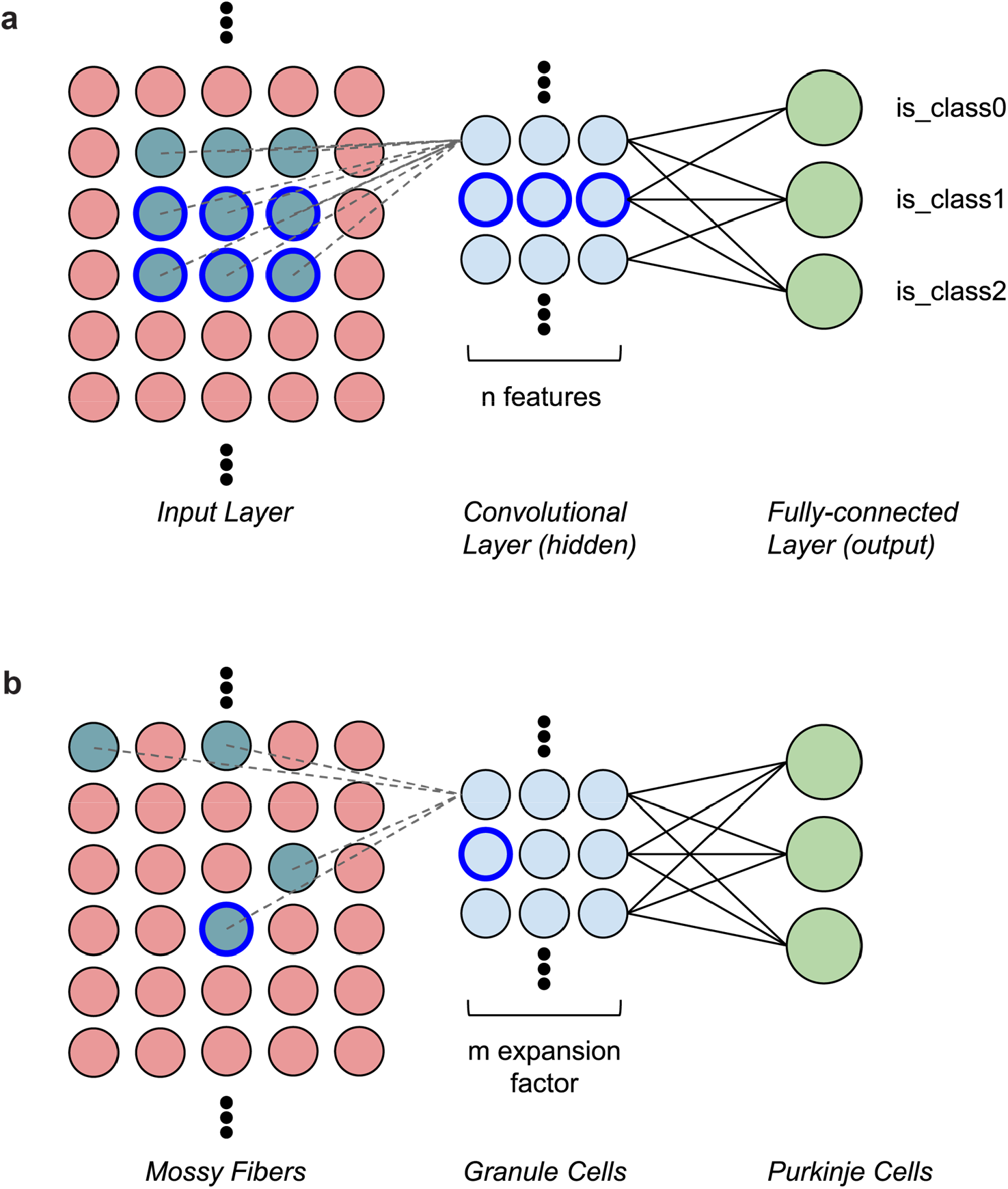
Similarity between a convolutional neural network and the cerebellar feedforward network. **a**, Diagram of a simple convolutional neural network with one convolutional layer (input→hidden) and one fully connected layer (hidden→output). The input (left) is made up of a single-channel 2D grid of neurons. The convolutional layer (middle) is made up of neurons each sampling a small local grid of the input (e.g., nine inputs when a 3×3 filter is used, cyan colored circles). This is notably different from a multi-layer perceptron network where the input and the hidden layer are fully connected - the convolution allowed an increase in features while decreasing computational cost. Due to the small field of view of each convolutional layer neuron, adjacent neurons share a significant amount of inputs with each other. To increase capacity of the hidden layer, the convolutional neurons can be replicated by *n* times (typically parameterized as *n* features). Finally, the output neurons (right) are fully connected with neurons in the preceding convolutional layer. This “fully-connected” layer is similar to the “linear” layers found in perceptron or linear/logistic regression networks. For a classification network, each label (class) is associated with a single binary output neuron for both training and inference. **b**, Diagram of the cerebellar feedforward network. Mossy fibers (MFs; left) can be considered a 2D grid of sensory and afferent command inputs typically of mixed modalities^53,54^. Granule cells (GrCs; middle) sample only ∼4 MF inputs each that are thought to be randomized, but with locality dependent on the length of the dendrites. Due to random connectivity, however, adjacent GrCs share few common inputs. The total number of GrCs is estimated to be hundreds of times more than the number of MFs (**Fig. 1b**), representing an expansion factor *m* of hundreds. Finally, Purkinje cells (PCs; right) - output neurons of the cerebellar cortex - are synapse onto by tens to hundreds of thousands of GrC axons that are passing by PC dendrites. PC axons converge in the deep cerebellar nuclei.

**Extended Data Figure 2.**
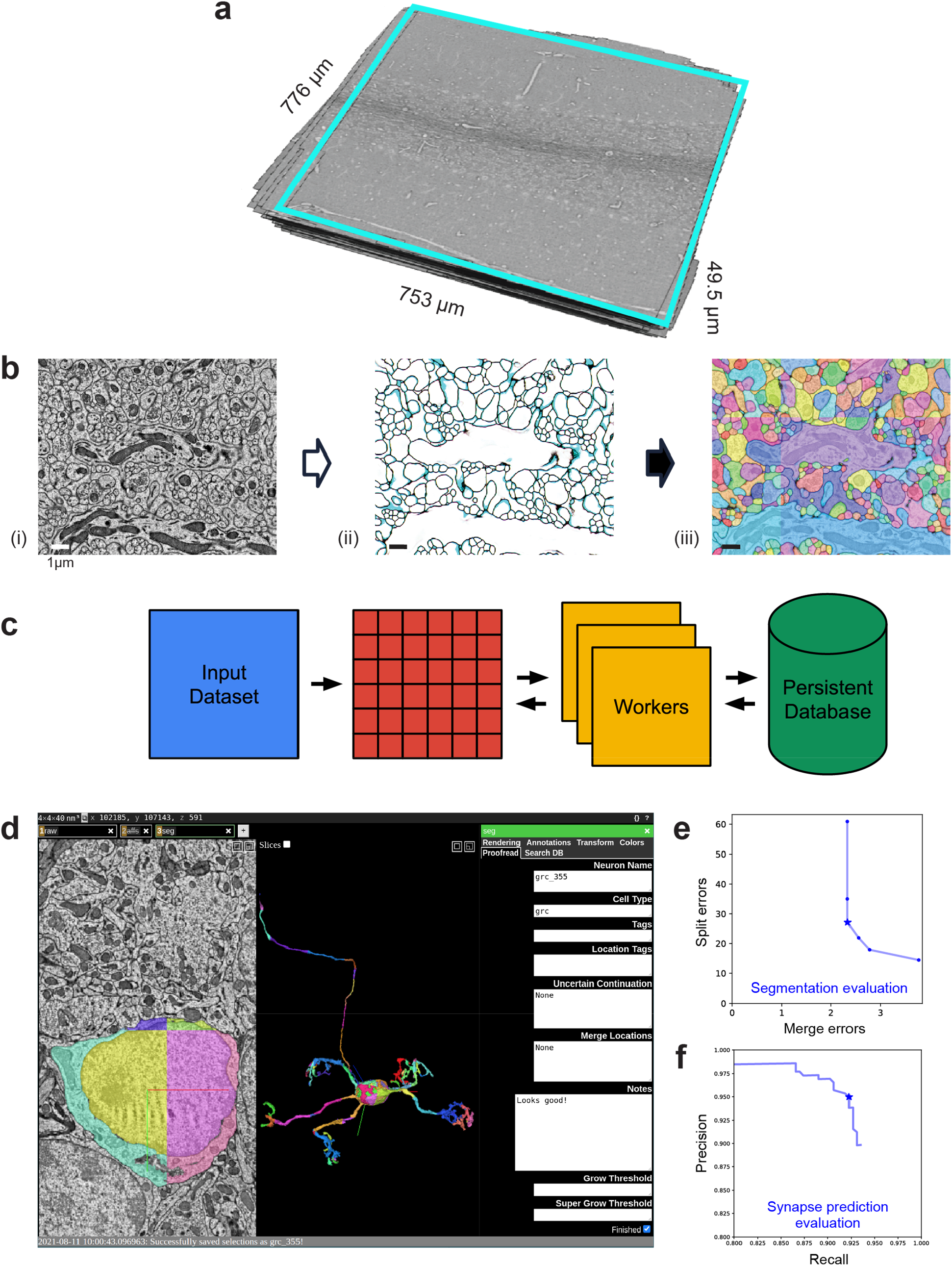
Automated segmentation and synapse prediction. **a**, Serial-section electron microscopy (EM) dataset from lobule V from the cyan boxed region in **Fig. 1d. b**, The 3D reconstruction segmentation pipeline. (i) EM image data, (ii) boundary affinities, and (iii) automated segmentation output. **c**, Parallelized volume processing using *daisy*. The input dataset is divided into small blocks, of which multiple workers can dynamically query for and work on. Block completion status and output data are efficiently stored into a persistent database or disks directly from the workers without going through the centralized scheduler process. **d**, Example view of targeted neuron reconstruction using merge-deferred segmentation (*MD-Seg*). Neurons are first segmented as small blocks, and inter-block merge decisions are deferred to proofreaders, hence the different colored segments constituting the shown neuron. The user interface is based on *Neuroglancer*, modified to provide the segment “grow” functionality, and to integrate an interface to the database keeping track of neuron name, cell type, completion status, notes, and which agglomeration threshold to use for “growing”, as well as searching for neurons based on different criteria and recoloring segments of a single neuron to a single color (“Search DB” and “Color” tabs, not shown). **e**, Automated segmentation evaluation; plot points denote agglomeration thresholds. Number of merge and split errors within 6 μm^3^ test volumes. We used a threshold (star) with 2.33 merges and 27 splits per 6 μm^3^ for proofreading. **f**, Automated synapse prediction evaluation; plot points denote connected component thresholds. Precision and recall curve for the synapse inference network. We achieved high synapse prediction accuracy with precision: 95.4% and recall: 92.2%, and an f-score: 93.8% (star).

**Extended Data Figure 3:**
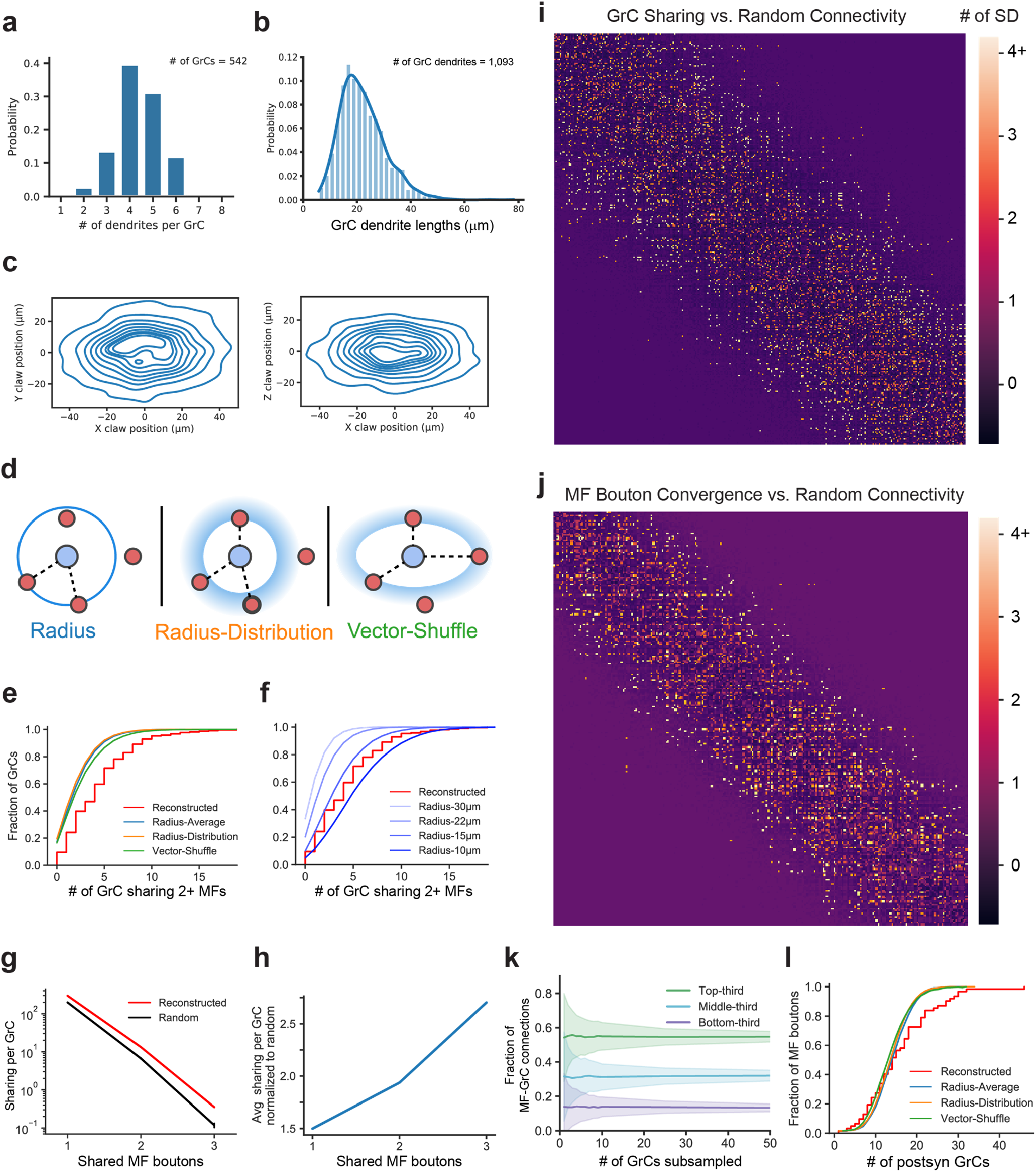
MF→GrC wiring, convergence, and null models. **a**, Distribution of the number of dendrites per GrC (n = 542). **b**, Distribution of GrC dendrite lengths (n = 1093). **c**, Anisotropic positioning of MF bouton→GrC inputs (claws), showing elongated distribution in the dorsal-ventral axis (X) relative to both anterior-posterior (Y) and medio-lateral (Z) axes. Contour lines represent 10% changes in the distribution increasing toward the center. **d**, MF→GrC random models used for comparison with the reconstructed connectivity. The *Radius* model randomly connects each GrC (blue circle) to a set of MF boutons (red circle, dashed lines indicate valid possible connections) closest to a fixed radius taken to be the average dendrite length, similar to models used in prior work^11–13,32^. The *Radius-Distribution* model is similar but draws radius randomly from the observed distribution in **b** for each MF→GrC connection. The *Vector-Shuffle* model draws a random vector from the anisotropic positioning of MF→GrC distribution in **c** to preserve the anisotropic distribution of MF→GrC connectivity (**Methods**). **e**, Same as **Fig. 2c**, but with random models from **d** added. **f**, Same as **e**, but with *Radius* models of different dendrite lengths. **g**, Average number of GrC pairs sharing 1, 2, or 3 common MF bouton inputs, comparing reconstructed against the *Radius* random connectivity model described in **d. h**, Average sharings of GrCs of the reconstructed network as in **g** but normalized to random networks. **i**, GrC input sharing relative to random connectivity. The matrix shows the degree of input sharing between GrCs (centermost n = 550, sorted by soma position rostrocaudally). The color scale for each cell in the matrix uses the z-score (reconstructed # of sharing minus random mean divided by the SD). **j**, MF bouton output convergence relative to random connectivity. The matrix shows the degree of output convergence between MF boutons (centermost n = 234, sorted by soma position rostrocaudally). The color scale uses the z-score as in **i. k**, Average fractional distribution of inputs to GrCs from different MF bouton types (categorized as the top-third, middle-third, and bottom third most connected boutons) as a function of GrC sampling size. GrCs were randomly subsampled to produce input composition distributions, with error shadings representing SD. **l**, Same as **Fig. 2e**, but with random models from **d** added.

**Extended Data Figure 4:**
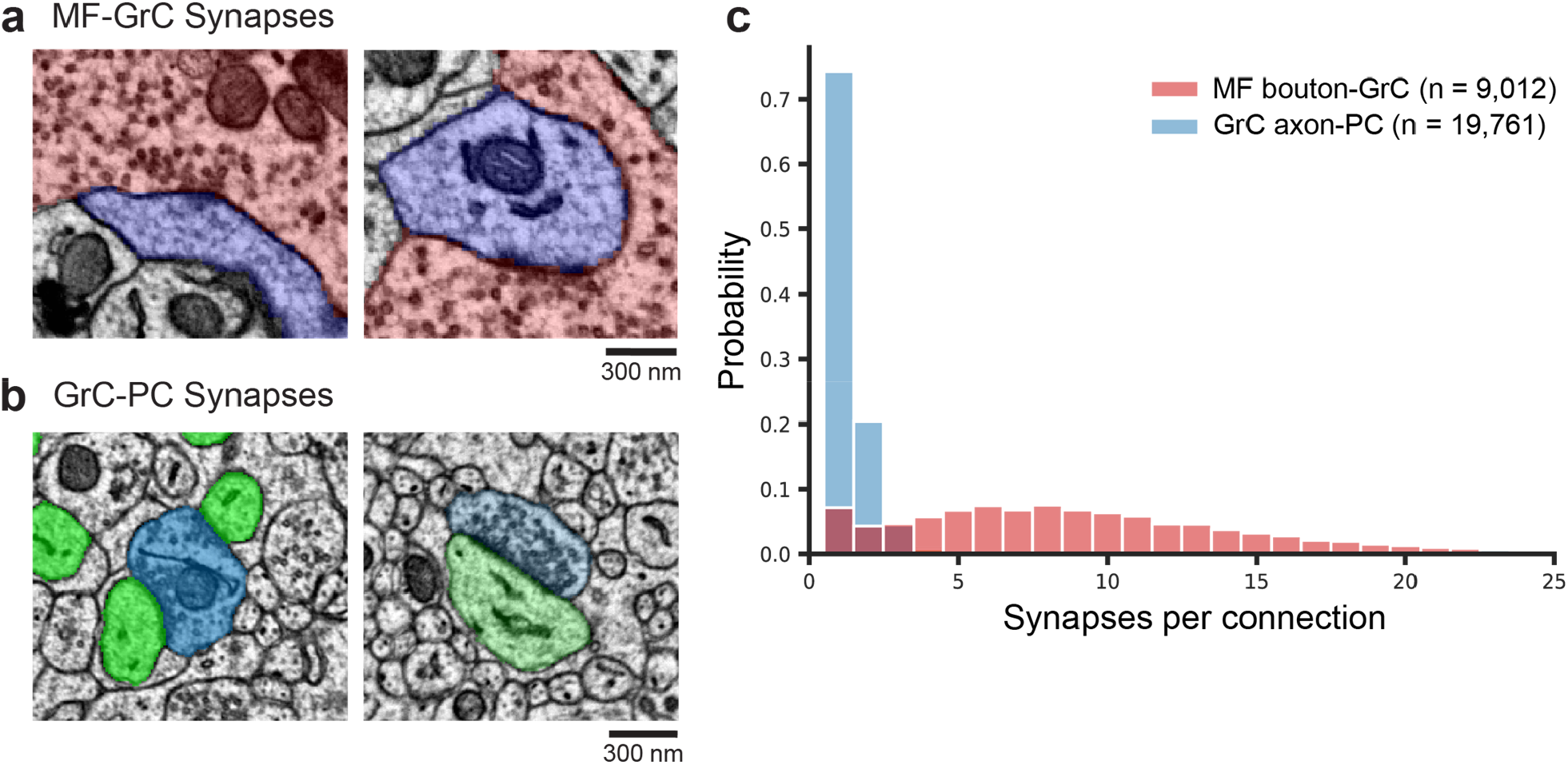
MF→GrC and GrC→PC synaptic connectivity. **a**, Examples of EM of MF (red) to GrC (blue) synapses. **b**, Examples of EM of GrC (blue) to PC (green) synapses. **c**, Distributions of the number of synapses per connection of the two synapse types. The median of MF→GrC is nine synapses, while GrC→PC is one. Over 97% of GrC→PC unitary connections have 1 to 2 synapses, though instances of 3 to 6 synapses per connection, while rare, do occur.

**Extended Data Figure 5:**
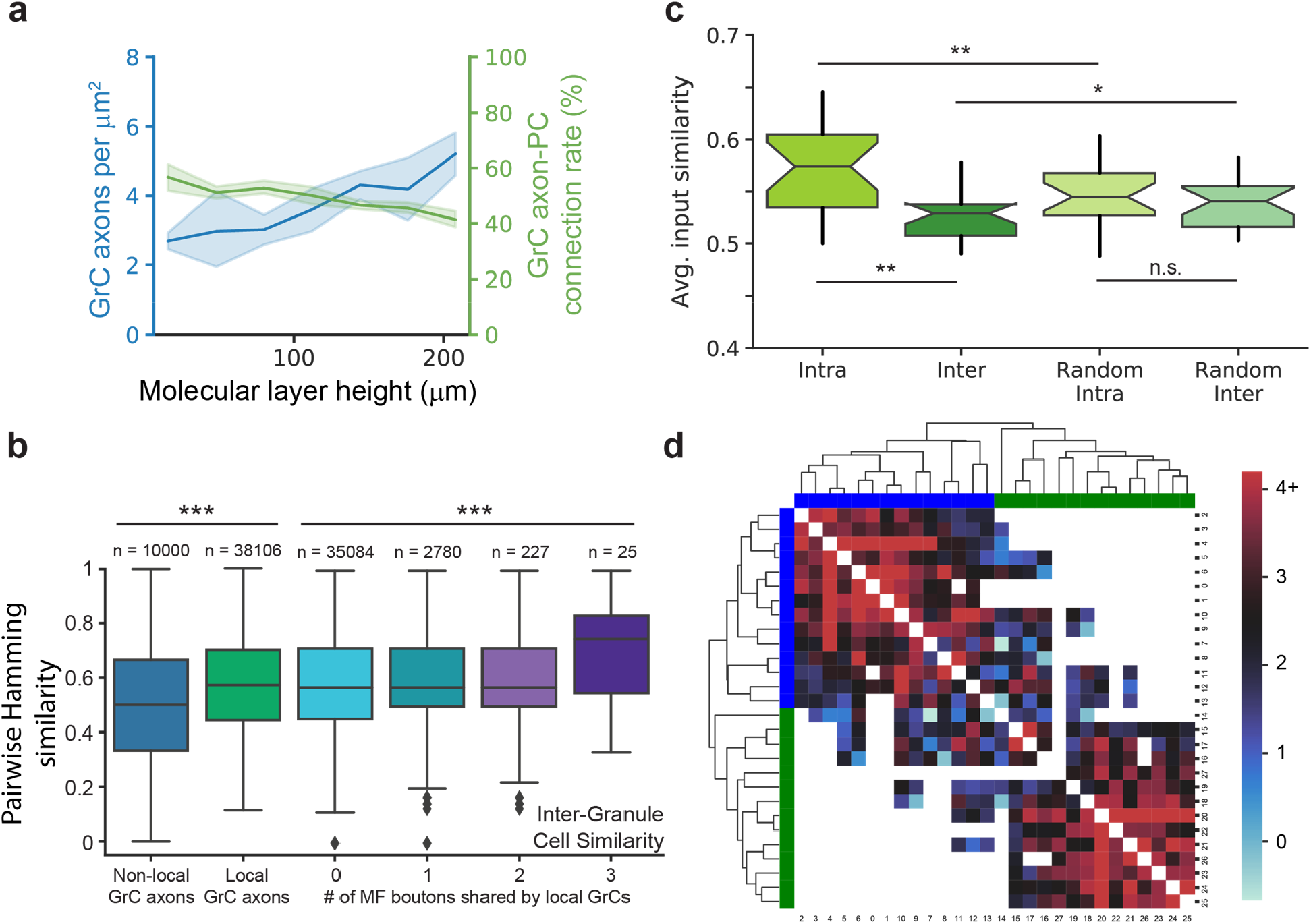
GrC→PC wiring and similarity of inputs to PCs. **a**, Plot of density GrC axons and GrC**→**PC connectivity rate as a function of height in the molecular layer between the pial surface and PC layer. Across molecular heights, the average axon density is 3.73 ± 1.23 per μm^2^ (mean ± SD), and the average connection rate is 49.12 ± 4.39% (mean ± SD). Using these numbers and the average area of PC dendrites, we calculated ∼125,000 GrC axons pass through the dendritic arbor of each reconstructed PC. At an average connectivity rate of 49%, only about 60,000 GrC axons were connected to each PC, 3-5× less than typically assumed in models of the cerebellar cortex^17,40,41^. **b**, Box plot of pairwise Hamming similarity between non-local GrC axons, and local GrC axons with different numbers of shared MF bouton inputs. Across local GrCs sharing 0, 1, 2, and 3 MF boutons, *p* = 0.0001137, Kruskal-Wallis H-test. 0-shared vs 1-shared *p* = 0.0132, 0-shared vs 3-shared *p* = 0.00797, 1-shared vs 3-shared *p* = 0.0186, 2-shared vs 3-shared *p* = 0.0309, other pairings *p* > 0.05, Dunn’s post hoc tests, Bonferroni corrected for multiple comparisons. **c**, Notched box plot of average Hamming similarity of inputs among intra-cluster PCs, inter-cluster PCs, random intra-cluster PCs, and random inter-cluster PCs from *k*-means clusters where *k* = 3. Intra vs. inter *p* = 0.000934, intra vs. random intra *p* = 0.000508, inter vs. random inter *p* = 0.0295, random intra vs random inter *p* = 0.575, Wilcoxon rank-sum tests, Bonferroni corrected for multiple comparisons. **d**, Input similarity matrix of PCs as in **Fig. 3f**, but grouped by hierarchical clustering yielded the same groups. The color bars denote the PCs that were colored blue or green in **Fig. 3e**, and the number labels denote the same PCs as ordered in **Fig. 3f**.

**Extended Data Figure 6:**
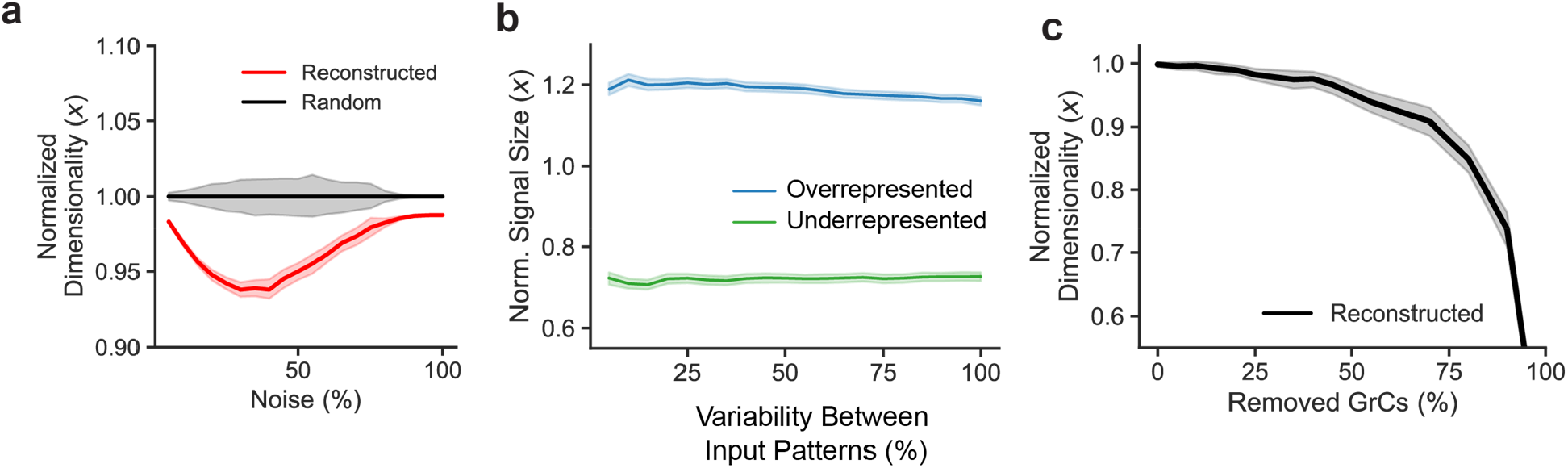
MF-GrC-PC simulations. **a**, Normalized dimensionality of the GrC population as a function of noise using spike frequency. Unlike the model in **Fig. 4c** where binary input patterns are used, this model represents the MF input patterns as continuous variables^13^ (spike frequency), with noise modeled as the variation of spiking frequency. The reconstructed network model exhibits lower dimensionality indicating less variability in the face of noise. **b**, Modeled learned signal size (as described in **Fig. 4e**) as a function of variability between MF input patterns, comparing overrepresented (top-third most connected boutons) vs underrepresented (bottom-third) MF boutons. Signal size from the reconstructed network is normalized by the random connectivity model for each population separately. **c**, Dimensionality of random inputs as a function of percentage of GrCs randomly removed, normalized to the dimensionality with 100% of the population.

## Supplementary Information for

**Supplementary Data 1: Reconstructed mossy fibers (MFs)**.

Example MFs reconstructed in the EM dataset. Zoomed-in region (top-left) from orange boxed region in the zoomed-out (bottom) view. An EM section (top-right) through the MF bouton (shaded purple) from the zoomed-in reconstruction (yellow box, top-left).

**Supplementary Data 2: Gallery of reconstructed granule cells (GrCs)**.

Example 3D reconstructions of GrCs.

**Supplementary Data 3: Gallery of reconstructed Purkinje cells (PCs)**.

Example 3D reconstructions of PCs.

**Supplementary Video 1. Fly-through of the cerebellar electron microscopy (EM) dataset**.

Zoomed-out (left) and zoomed-in (right) views of aligned EM image data.

**Supplementary Video 2. Reconstructed granule cells (GrCs)**.

3D volumetric reconstructions of example GrCs.

**Supplementary Video 3. Reconstructed Purkinje cells (PCs)**.

3D volumetric reconstructions of example PCs.

