## Supplementary figures and images for "Structured connectivity in the cerebellum enables noise-resilient pattern separation"

### Supplementary Data 1

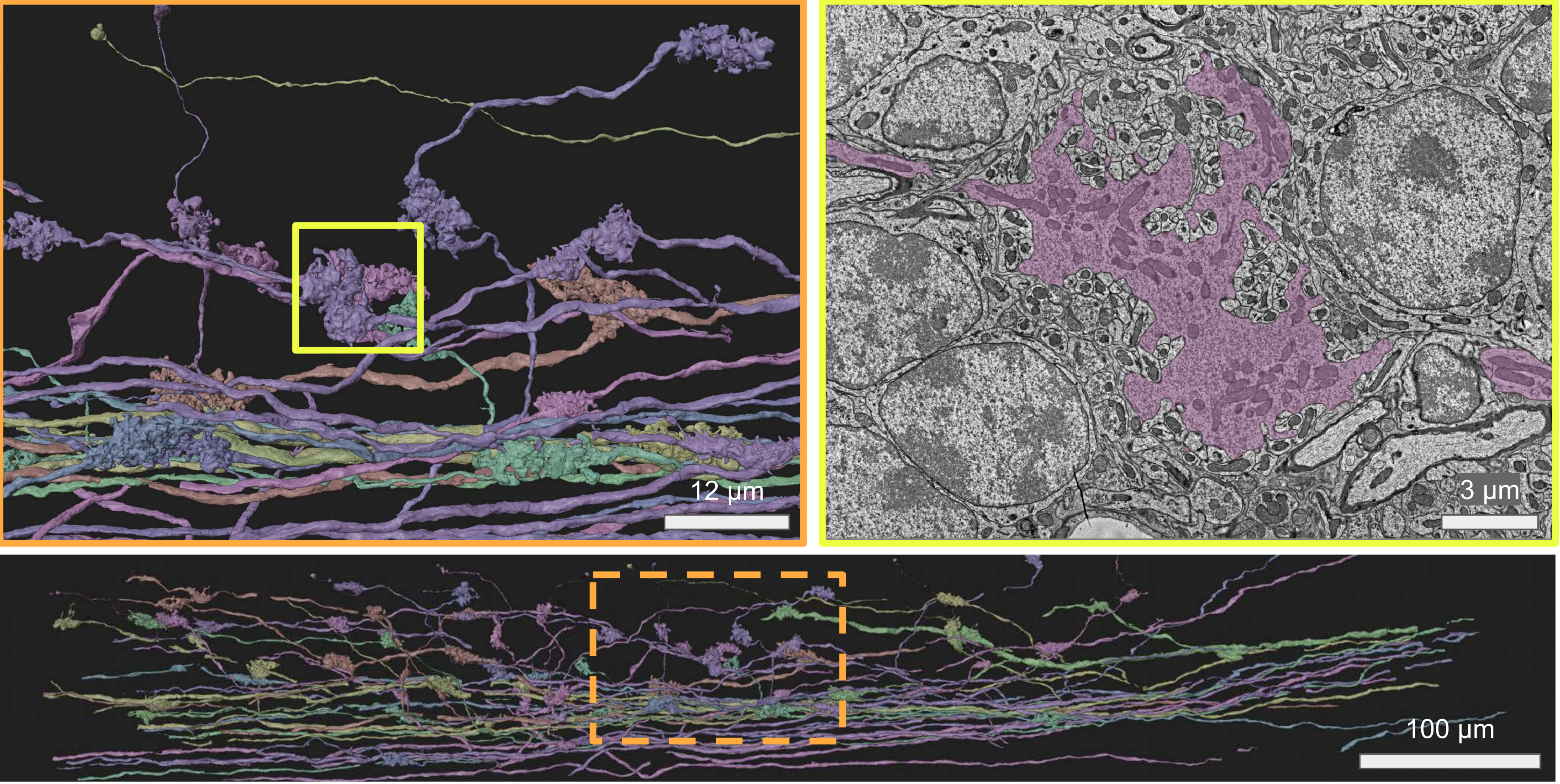

### Supplementary Data 2

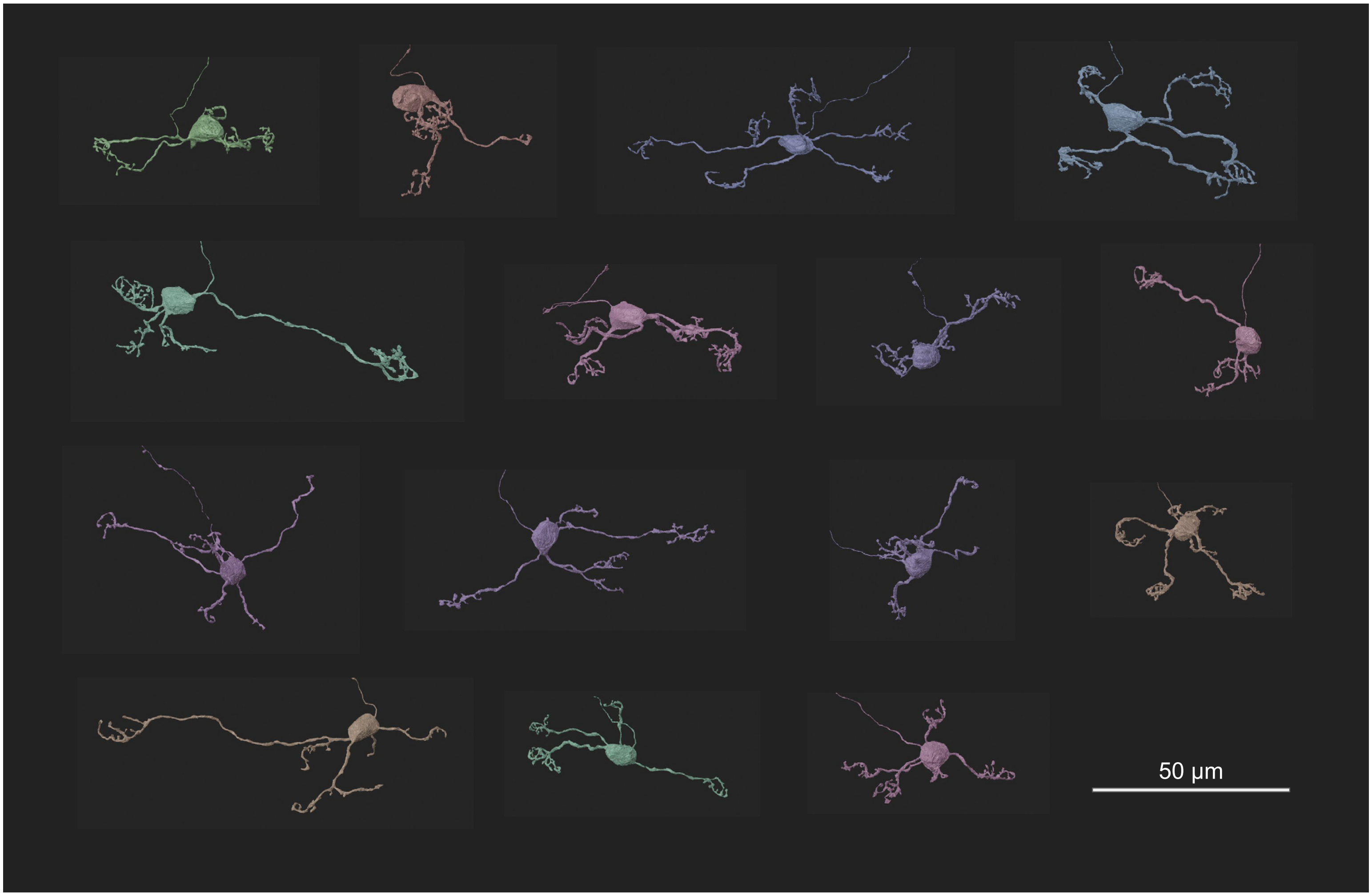

### Supplementary Data 3

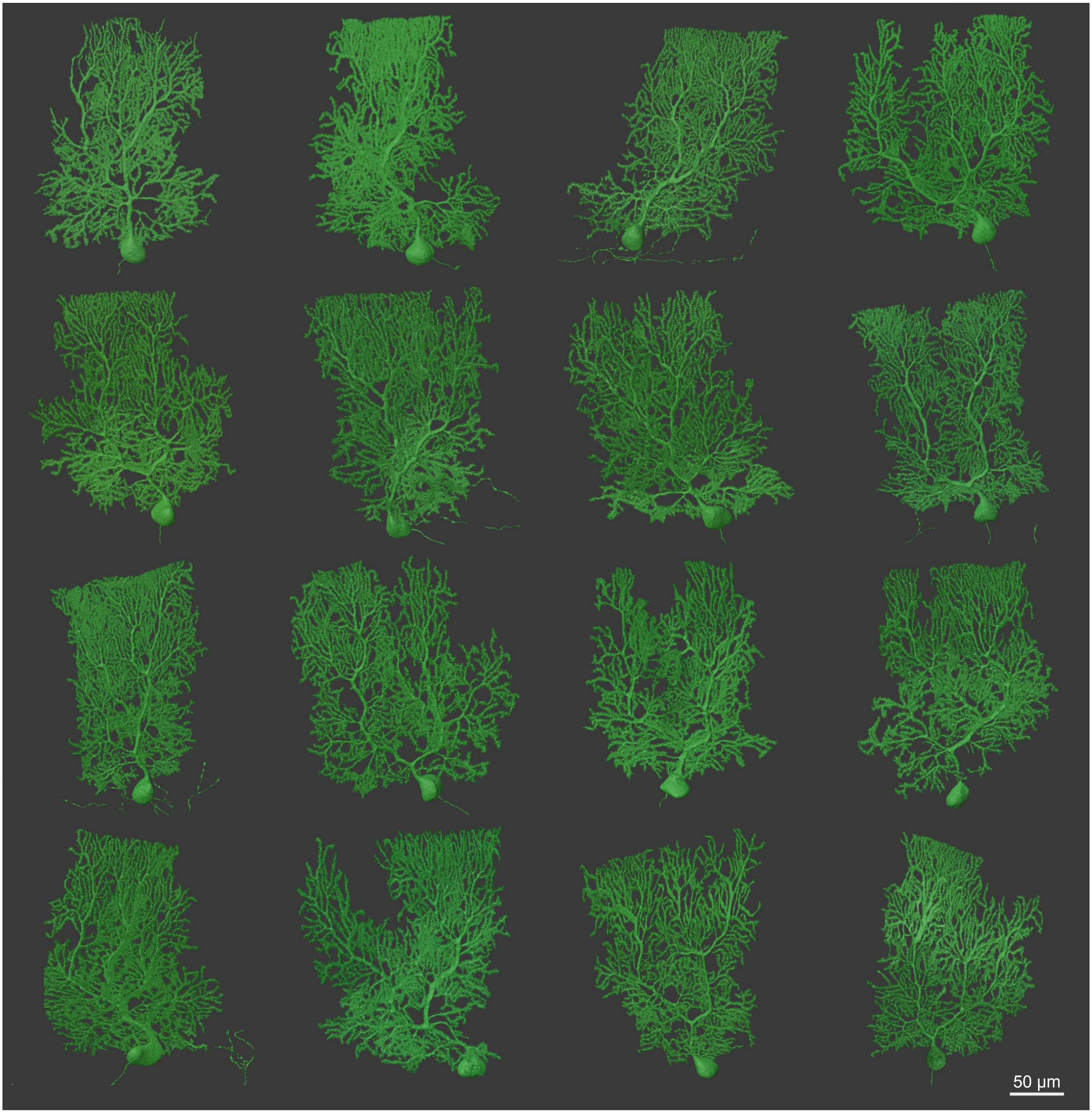
